# Unveiling unique microbial nitrogen cycling and novel nitrification drivers in coastal Antarctica

**DOI:** 10.1101/2023.11.19.566674

**Authors:** Ping Han, Xiufeng Tang, Hanna Koch, Xiyang Dong, Lijun Hou, Danhe Wang, Qian Zhao, Zhe Li, Min Liu, Sebastian Lücker, Guitao Shi

## Abstract

Although microbial nitrogen (N) cycling plays a pivotal role in Antarctic ecosystems, its underlying mechanisms are largely uncharted. In this study, we unravel the biological origin of nitrate via triple oxygen isotopic composition analysis and systematically profile functional N-cycling genes within soil and lake sediment samples from the ice-free areas of East Antarctica. We successfully reconstruct 1,968 metagenome-assembled genomes (MAGs) spanning 29 microbial phyla, enabling the analysis of the presence or absence of 52 diverse metabolic marker genes. Consistent with quantitative data, our metagenomic analyses confirm the active processes of microbial nitrogen fixation, nitrification, and denitrification. We find no detectable anaerobic ammonium oxidation (anammox) processes, underscoring a unique microbial N-cycling dynamic in the region. Notably, we identify the predominance of complete ammonia-oxidizing (comammox) *Nitrospira*, a recently discovered bacterial guild capable of performing the entire nitrification process within a single organism. Further genomic investigations reveal their adaptive strategies in the Antarctic environment. These strategies likely involve the synthesis of trehalose to counteract cold stress, high substrate affinity to efficiently utilize available resources, and alternative metabolic pathways to adapt to nutrient-scarce conditions. Their significant role in the nitrification process is validated through ^13^C-DNA-based stable isotope probing (DNA-SIP). This research provides a comprehensive illustration of nitrification’s crucial contribution to the nitrogen budget in coastal Antarctica, highlighting comammox *Nitrospira* clade B as a novel nitrifying agent and shedding new light on the complex biogeochemical processes of nitrogen cycling in coastal Antarctica.

## Introduction

Antarctica, a region largely insulated from anthropogenic nitrogen (N) deposition, exhibits notably low N concentrations^1^. These diminished N levels play an integral role in sustaining ecological equilibrium in the continent’s barren, ice-free terrains^2, 3^. Consequently, elucidating the intricacies of microbial nitrogen cycling pathways in Antarctic ecosystems—including diazotrophy (N-fixation), nitrification, anaerobic ammonium oxidation (anammox) and denitrification—is paramount^4^. Diazotrophic Cyanobacteria are the primary agents of biologically accessible ammonium (NH_4_^+^) synthesis in these environments^5^. Alongside NH_4_^+^, nitrate (NO_3_^−^) constitutes a significant inorganic nitrogen pool in polar biomes^6, 7^. The genesis of NO_3_ ^−^ in Antarctic ecosystems can be attributed to both abiotic atmospheric deposition, particularly within mineral-rich Antarctic soils^8^, and to biotic nitrification processes. Still, the discrete contributions of these sources to the coastal Antarctic NO_3_^−^ reservoir remain inadequately characterized, necessitating further investigation.

Nitrification plays a pivotal role in the global biogeochemical nitrogen cycle and contributes significantly to the emissions of nitrous oxide (N_2_O), a potent greenhouse gas. This process encompasses the oxidation of ammonia (NH_3_) to nitrate (NO_3_^−^) via the intermediate nitrite (NO_2_^−^). Traditionally, chemolithoautotrophic nitrification has been attributed to two distinct microbial guilds: ammonia-oxidizing bacteria (AOB) and archaea (AOA), responsible for converting NH_3_ to NO_2_^−^, and nitrite-oxidizing bacteria (NOB), which facilitate the subsequent oxidation of NO_2_^−^ to NO_3_^−^. Beyond these canonical nitrifiers, the recently identified complete ammonia oxidizers (comammox) within the genus *Nitrospira*, specifically lineage II, perform full nitrification as a singular ^9, 10^. These organisms are increasingly recognized for their significant role in nitrification across diverse ecosystems, including both engineered systems^11^ and natural environments^12^. The contribution of comammox bacteria to nitrification and subsequent NO_3_^−^ production in Antarctic ecosystems, however, remains an enigma.

Phylogenetically, comammox *Nitrospira* bifurcate into clades A and B^9^, distinguished by their respective ammonia monooxygenase genes (*amoA*). Physiological assessments of clade A comammox *Nitrospira* have revealed an exceptionally high ammonia affinity, suggestive of an oligotrophic niche adaptation ^13, 14^. In stark contrast, clade B comammox *Nitrospira*, despite their widespread distribution in various habitats, including those characterized by low temperatures such as the Tibetan Plateau^15, 16^ and Arctic permafrost^17^, have yet to be cultured for physiological scrutiny.

This study aims to unravel the complexities of the microbial nitrogen cycle in the ice-free regions of East Antarctica, identifying the primary sources of NO_3_^−^ and examining the potential roles of novel microbes in the nitrification process. Our research is strategically centered on the Larsemann Hills (LH)^18^, a vast ice-free rocky landscape in East Antarctica, dotted with over a hundred oligotrophic lakes shrouded in ice. Through a comprehensive approach, we discover that (i) the origin of NO_3_^−^ is primarily the biological nitrification process; (ii) the microbial nitrogen cycle is distinct, encompassing most microbial N-cycling processes except for the anammox pathway; and (iii) the comammox *Nitrospira* clade B serves as a novel, abundant, and active driver of nitrification, possessing unique survival strategies against the cold and oligotrophic conditions of the coastal Antarctic environment.

## Results and Discussion

### Isotopic signatures indicate the biological origin of nitrate

Depending on how NO_3_^−^ is produced, the composition of its oxygen isotopes differs, allowing differentiation between abiotic and biotic sources^19^. For biologically produced NO_3_^−^, one oxygen atom (O) is expected from atmospheric oxygen (O_2_) and two from the surrounding water (H_2_O)^19^. Biologically produced NO_3_^−^ would have a δ^18^O of ∼0.6‰, since δ^18^O of atmospheric O_2_ is 23.9‰^20^ and the measured δ^18^O of LH lake water is –12.7±1.5‰ (Supplementary Table 1). Atmospheric NO_3_^−^ in Antarctica is mainly produced by the reaction of nitrogen oxides (NO_x_), ozone (O_3_), and hydroxyl radicals (OH·)^21^. In addition, while biologically produced NO_3_^−^ has a Δ^17^O of 0‰, which is identical to that of O_2_ and H_2_O, atmospheric NO_3_^−^ is usually characterized by high oxygen isotopic ratios^21, 22^ and can reach Δ^17^O ≥35‰ in the atmosphere in Antarctica^22^.

The mean δ^18^O of NO_3_^−^ in lake sediments from LH is 4.6‰. The annual mean δ^18^O of NO_3_^−^ in snow and the atmosphere at the coastal Zhongshan station, situated in LH, is approximately 73‰^21^. From these, the contributions of atmospheric deposition and biological production to sedimentary NO_3_^−^ pools can be quantified by isotope mass balancing, indicating that the nitrification process constitutes 93% of the NO_3_^−^ in sediments. In addition, limited data of sediment NO_3_^−^ Δ^17^O produced a mean of approximately 1.3‰, which strengthens the observed >90% contribution of nitrification considering the very high Δ^17^O of atmospheric NO_3_^−^ at Zhongshan station (approximately 35‰). Similarly, the isotope mass balance of δ^18^O and Δ^17^O suggested ∼96% of soil NO_3_^−^ in the two study areas is from nitrification. Thus, we demonstrated here that sediment and soil NO_3_^−^ predominantly originated from biological production in and nearby the investigated lakes.

### Unique microbial nitrogen cycle in coastal Antarctica

The primary production of NO_3_^−^ is largely attributed to the microbial nitrification process, rather than atmospheric precipitation. Consequently, the origin of NH_4_^+^, the substrate for nitrification, can be traced back to biological N-fixation. Based on the qPCR-based quantification of functional N-cycling genes (Fig. 1), we noticed a comparable amount of N-fixation (*nifH,* encoding nitrogenase) to nitrification genes (*amoA* and *nxrB*, encoding ammonia monooxygenase and nitrite oxidoreductase, respectively), with quantities ranging from 10^2-^10^4^ copies ng^−1^ DNA (Fig. 1). The relatively low quantity of the gene encoding hydroxylamine dehydrogenase (*hao*) can be attributed to the low coverage of the applied primers and the absence of the bacterial *hao* gene in the genomes of AOA^23^, which are of notable abundance in the studied samples.

**Fig. 1.**
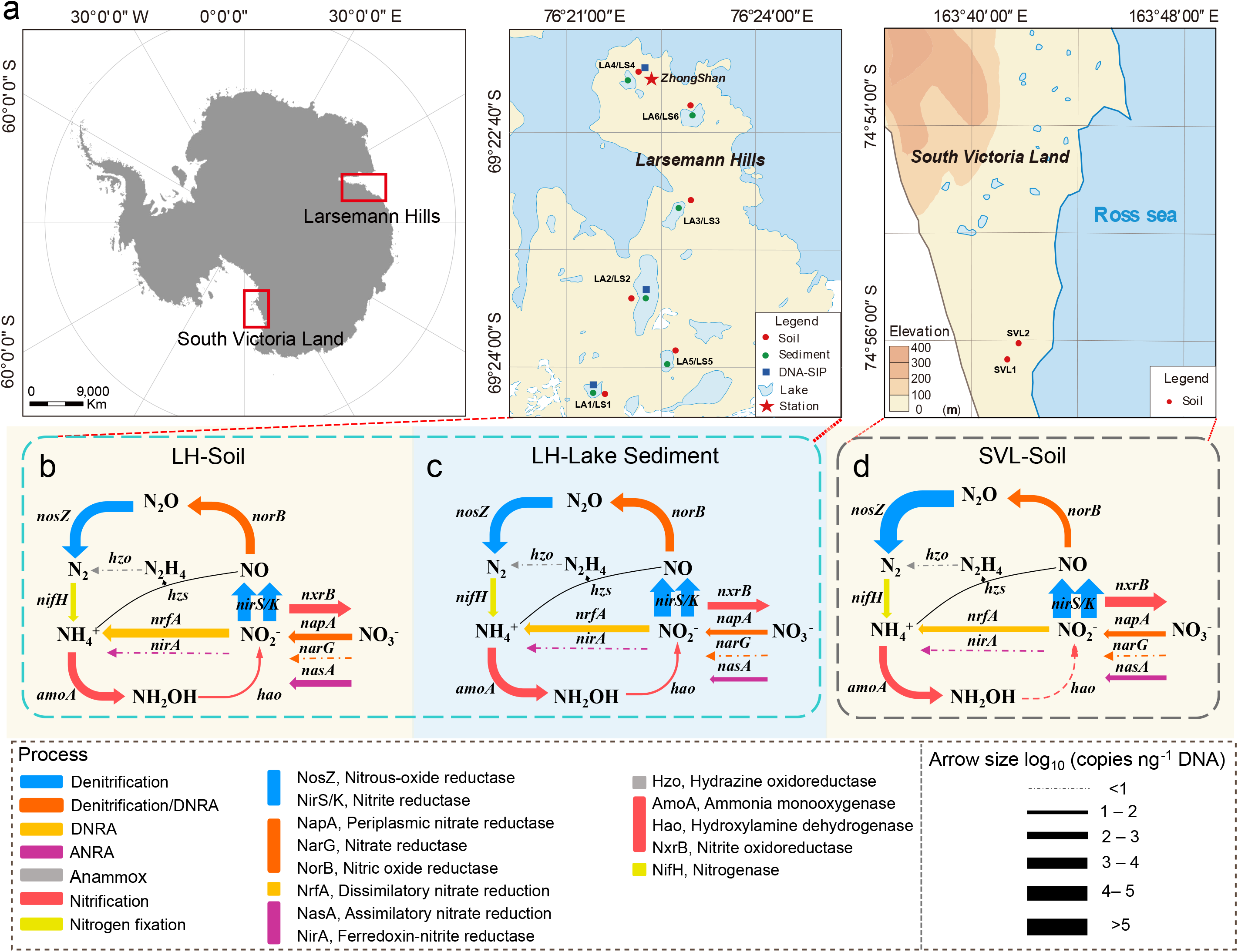
Overview of sampling sites, sample types, and quantities of functional nitrogen cycling genes in terrestrial Antarctica. **(a)** A photograph illustrating the location of the Larsemann Hills (LH) and South Victoria Land (SVL) (left) in Antarctica, locations of studied surface soil and lake sediments in LH (center), and locations of surface soil samples SVL (right). Red and green circles denote soil and sediment samples, respectively. **(b-d)** Abundancec of functional nitrogen cycle genes in the LH soils **(b)**, LH lake sediments **(c)**, and SVL soils **(d)**. The studied functional microbial nitrogen cycle genes encompass nitrogen fixation (*nifH*), nitrification (*amoA*, *hao*, and *nxrB*), denitrification (*napA*, *narG*, *nirS/K*, *norB*, and *nosZ*); dissimilatory nitrate reduction to ammonium (DNRA; nrfA), assimilatory nitrite reduction (ANR; *nasA* and *nirA*), and anaerobic ammonium oxidation (anammox; *hzo*). The relative abundance of the associated functional genes is indicated by the line thickness.

In addition to N-fixation and nitrification processes, we also observed significant activity of denitrifier, as indicated by the high abundances of genes encoding the sequential reduction of NO_3_^−^ (*nar, nap*), NO_2_^−^ (*nir*), nitric oxide (NO, *nor*), and N_2_O (*nos*) (Fig. 1). The relatively higher abundances of *nirS* and *nirK* genes can likely be attributed to their presence not only in denitrifiers but also in nitrifiers. Interestingly, we found substantial quantities of the functional gene for dissimilatory nitrate reduction to ammonium (DNRA, *nrfA*) and assimilatory nitrite reduction (ANR, *nasA*) (Fig. 1). The potential for DNRA and ANR is widespread among phylogenetically diverse microorganisms and these processes ensure the retention of inorganic nitrogen in an ecosystem.

Strikingly, we detected minuscule amount of the *hzo* gene (encoding hydrazine oxidoreductase), a biomarker for the anammox process, in all tested sediment and soil samples. Anammox bacteria catalyze the anaerobic oxidation of NH_4_^+^ using NO_2_^−^ as electron acceptor, producing N_2_ as final product. These bacteria were first identified in wastewater treatment systems^24^ and were subsequently discovered in various environments, including marine^25^, coastal^26^, terrestrial^27^ and engineering systems^28^. The absence of anammox functional markers in coastal Antarctica suggests unique microbial nitrogen cycling properties in this remote region.

### Highly diversity of microbiomes and their roles in microbial N-cycling processes

From an extensive dataset exceeding 280 gigabases of sequencing data, we managed to reconstruct a dereplicated collection of 724 high-quality and 1244 medium-quality^29^ metagenome-assembled genomes (MAGs) (Supplementary Data 1). The recovered genomes encompass 29 distinct phyla (Supplementary Data 2), marking, to the best of our knowledge, the most comprehensive inventory of Antarctica coastal soil and sediment genomes to date. On average, we obtained 50 to 60 MAGs per sample site studied (Fig. 2a). The most abundant phyla included *Actinobacteriota*, *Pseudomonadota*, *Bacteroidota*, *Chloroflexota*, *Verrucomicrobiota*, *Acidobacteriota*, *Patescibacteria*, *Planctomycetota*, and *Gemmatimonadota* (Fig. 2b, Supplementary Data 1), aligning with findings from other Antarctic surveys ^5, 30, 31^. *Cyanobacteriota* also featured among the top ten most abundant MAGs (Supplementary Data 1), a deviation from the Mackay Glacier Region, where they were largely absent in most soil samples^31^. We identified only five archaeal MAGs within the Thermoproteota phylum, all of which were further classified as ammonia-oxidizing archaea (AOA) within Group I.1b.

**Fig. 2.**
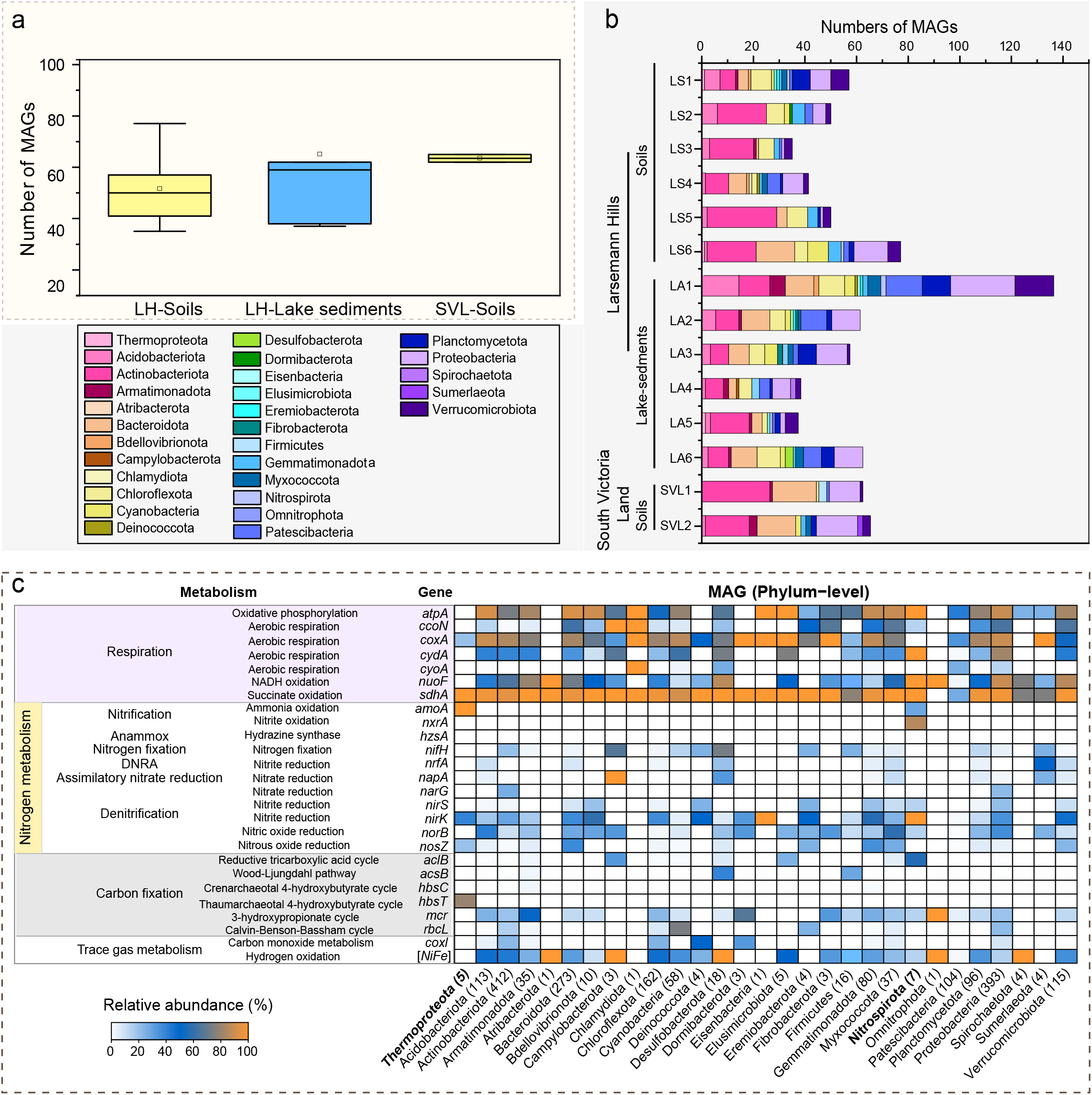
Community composition and relative abundance of selected functional genes derived from metagenomic analyses of various sediments and soils from the coastal East Antarctic regions. **(a)** Number of metagenome-assembled genomes (MAGs) retrieved from soil and sediment samples collected from the Larsemann Hills (LH), as well as soil from South Victoria Land (SVL). **(b)** Composition of the microbial community across all analyzed samples. **(c)** Distribution of selected key functional genes in the obtained MAGs, including those involved in respiration, nitrogen cycling, carbon fixation, and trace gas metabolism. The adjacent heatmap displays the distribution of these genes across a set of 1968 MAGs, comprising 29 phyla.

To comprehend the metabolic strategies that sustain the abundant bacterial life in these extremely nutrient-poor sediments and soils, we examined the distribution and affiliation of 52 marker genes conserved across different energy conservation and carbon acquisition pathways in the MAGs we retrieved. As predicted, genes for aerobic organotrophic respiration were encoded by nearly all community members (Fig. 2c, Supplementary Fig.1). Consistent with observations from the Mackay Glacier Region^31^, a significant number of the MAGs appear to fix carbon via the Calvin-Benson-Bassham (CBB) cycle or the 3-hydroxypropionate cycle (Fig. 2c). These processes provide a pathway for biomass generation that is independent of photoautotrophy, a function primarily performed by *Cyanobacteriota* (Supplementary Fig.1). Genomic analysis revealed that the most abundant and widespread community members encoded trace gas oxidation genes. In a pattern similar to previous discoveries ^31, 32^, carbon monoxide (CO) dehydrogenases (CoxL) were exclusive to *Actinobacteriota* and *Chloroflexota* (Fig. 2c, Supplementary Data 3). Interestingly, uptake hydrogenases were encoded by MAGs from *Acidobacteriota*, *Chloroflexota*, and *Verrucomicrobiota* (Fig. 2c, Supplementary Data 3), which aligns with observations from temperate soil where *Acidobacteriota* are known to be active atmospheric H_2_ consumers^33^, albeit with slight differences.

In the realm of N-metabolism, a comprehensive array of functional genes was identified within the MAGs we analyzed (Fig. 2c). Nitrogen fixation appears to be predominantly carried out by taxa within *Acidobacteriota*, *Cyanobacteriota*, Desulfobacterota, Myxococcota, and Verrucomicrobiota (Fig. 2c). Consistent with the quantitative analyses, the *hzs* gene, indicative of the anammox process, was not present in the retrieved MAGs. While a total of 96 *Planctomycetota* MAGs were obtained, subsequent verification through taxonomic classification to check for affiliation with the *Brocadiaceae* confirmed that none of these MAGs belong to anammox bacteria (Supplementary Data 2). The absence of anammox process may be attributed to the discrepancy between the optimal temperature range of 12–17°C, which is characteristic of anammox process found in similarly cold Arctic fjord sediments ^34^, suggesting a limited tolerance to low temperatures. This assertion is further reinforced by the absence of anammox bacteria in the McMurdo Dry Valleys (an Antarctic desert) ^35^ and in Arctic regions ^36^ as well. The pervasive distribution of denitrification genes across nearly all phyla underscores the robust and widespread microbial denitrification activity in Antarctica, corroborating previous research that highlights the ubiquity of cold-adapted denitrifiers across diverse Antarctic ecosystems^37^. Regarding the nitrification process, the marker genes-*amoA* (associated with ammonia oxidation) and *nxrB* (linked to nitrite oxidation) were found exclusively in the *Thermoproteota* and *Nitrospirota* phyla (Fig. 2c). Although Antarctic soil nitrification was previously reported back in 1997^38^, attributed to AOB genera *Nitrosospira* and *Nitrosomonas*, as well as AOA within group I.1b ^39–42^, no investigation has yet been conducted into the presence and function of the recently discovered comammox bacteria within the *Nitrospira* genus ^9, 10^. Here, we present data showing the prevalence of the comammox *Nitrospira amoA* gene in Antarctica lake sediments and soils, strongly suggesting the activity of comammox *Nitrospira* in this region.

### Abundant and novel nitrification drivers in coastal Antarctica

The quantitative evaluation of nitrification-related functional genes, including *amoA* and *nxrB* (Supplementary Fig. 2), established that AOA and comammox bacteria were the predominant nitrifiers in the analyzed soils and sediments (Supplementary Fig. 3). Our metagenomic analysis corroborates these findings, having identified only AOA and *Nitrospira* MAGs among the various nitrifying groups (Fig. 2c), suggesting their relatively high abundances (Supplementary Table 3). Furthermore, a more detailed investigation into the abundance and community structure of all identified nitrifying organisms using amplicon sequencing and subsequent phylogenetic assessment indicated an unexpected dominance of clade B comammox *Nitrospira* in this environment (Supplementary Results and Discussion).

Among the de-replicated MAGs, five were classified as Group I.1b-AOA within the genera *Nitrosocosmicus* and *Nitrospharea* of Thermoproteota phylum (Supplementary Data 2). This categorization stemmed from phylogenomic assessments and average nucleotide identity (ANI) analyses (Supplementary Fig. 4). These findings are in agreement with the data obtained from amplicon sequencing, as discussed in the Supplementary Results and Discussion section.). Together with their previous detection in Arctic soils ^43, 44^, this suggests a capacity of *Nitrosocosmicus*-like AOA to thrive in cold and nutrient-deficient conditions.

Of the seven *Nitrospirota* MAGs analyzed, four were verified as comammox *Nitrospira*, and the remaining three belonged to the exclusively nitrite-oxidizing lineages II and IV of *Nitrospira*. This classification was substantiated by phylogenomic and ANI analyses (Fig. 3a and 3b, Supplementary Fig. 5) and these results concur with the phylogenetic assessment based on the *amoA* and *nxrB* gene of *Nitrospira* (Supplementary Fig. 8 & Fig. 9). The notably high abundance of the AOA and these *Nitrospira* MAGs in the soil and sediment metagenomes studied (reaching up to 0.173%, Supplementary Table 3) further emphasizes their significant roles in nitrification within these ecosystems.

**Fig. 3.**
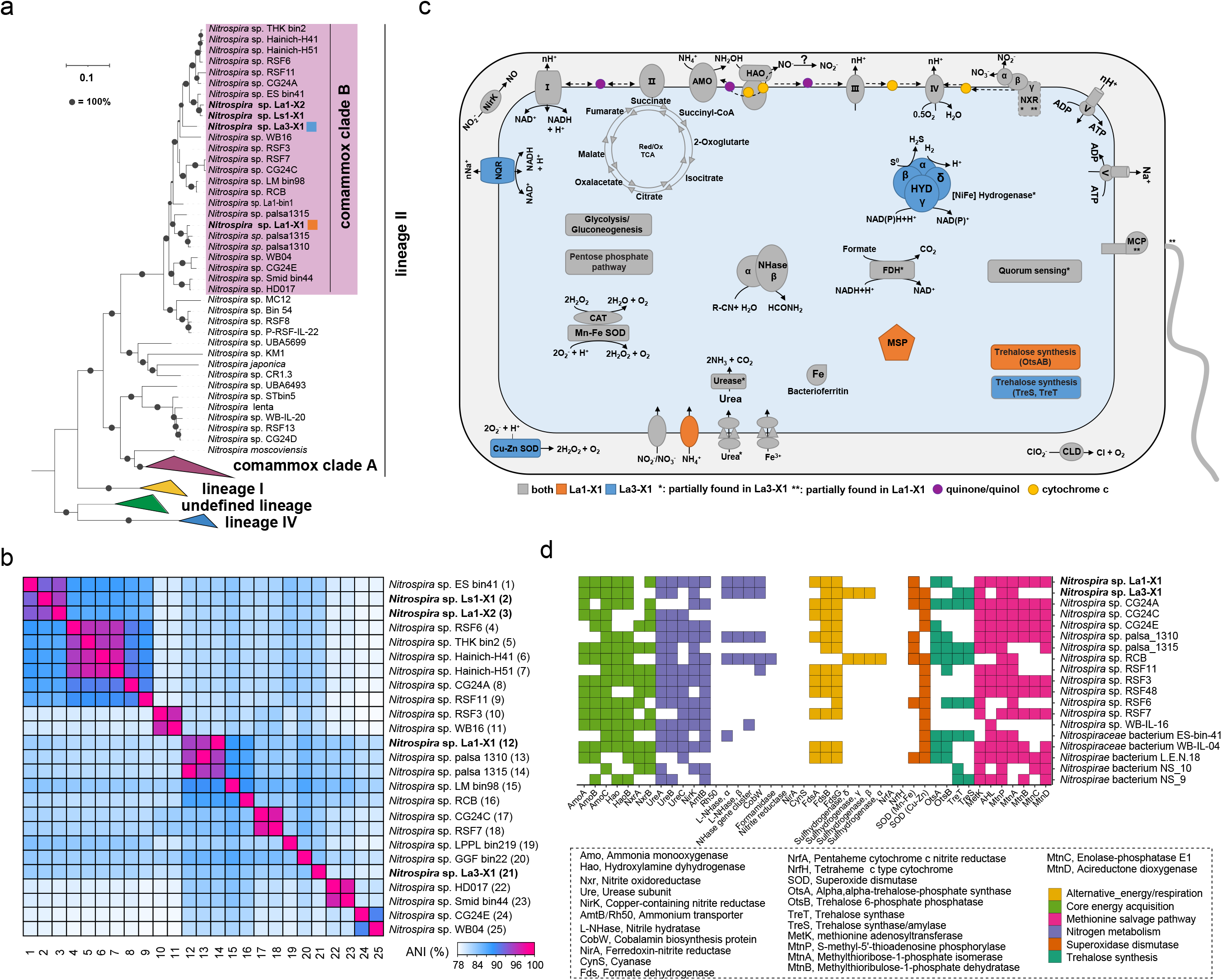
Phylogenomic and metabolic analyses of the comammox *Nitrospira* metagenome-assembled genomes (MAGs), providing insights into their survival strategies in coastal East Antarctica. **(a)** Maximum likelihood phylogenomic tree of the genus *Nitrospira*, constructed using a concatenated alignment of 91 single-copy core genes. **(b)** Average nucleotide identity (ANI) analysis of available clade B comammox *Nitrospira* genomes in the NCBI database. **(c)** Comparative cellular metabolic diagram for MAGs *Nitrospira* sp. La1-X1 and *Nitrospira* sp. La3-X1. Selected features are depicted, including core pathways of chemolithoautotrophic ammonia and nitrite oxidation, alternative energy metabolism, and potential detoxification pathways. Colors denote the presence of specific genes in La1-X1 (orange), La3-X1 (blue), or both (grey). Asterisks represent incomplete features. Key enzymes and pathways are abbreviated as follows: AMO, ammonia monooxygenase; CAT, catalase; CLD, chlorite dismutase; FDH, formate dehydrogenase; HAO, hydroxylamine dehydrogenase; HYD, 3b [NiFe] hydrogenase; MCP, methyl-accepting protein; MSP, methionine salvage pathway; NHase, nitrile hydratase; NirK, Cu-dependent nitrite reductase; NQR, Na^+^-translocating NADH: ubiquinone oxidoreductase; NXR, nitrite oxidoreductase; SOD, superoxide dismutase. Enzyme complexes of the respiratory chains are labeled using Roman numerals. **(d)** Distribution of key metabolic features involved in nitrogen and alternative energy metabolism, the methionine salvage pathway, superoxide dismutase, and trehalose synthesis. In total, 19 *Nitrospira* genomes, including the two high-quality clade B comammox genomes (La1-X1 and La3-X1) obtained in this study, were analyzed. Features shown in white were not detected.

Two high-quality comammox MAGs, *Nitrospira* sp. La1-X1 (hereafter referred to as “La1”) and *Nitrospira* sp. La3-X1 (hereafter referred to as “La3”), had genome sizes of 4.26 Mb and 3.78 Mb, respectively (Supplementary Table 3). The *amoA* gene sequences from La1 and La3 align closely with the dominant *amoA* OTUs and with previously published sequences from clade B comammox (Fig. 4c and Supplementary Fig. 8). Consistent with the *amoA*-based phylogenetic findings, phylogenomic analysis also demonstrates a distinct clustering of these comammox MAGs within clade B (Fig. 3a). For La1, the ANI was highest with MAG *Nitrospira* sp. palsa1310, which was identified from Arctic permafrost soil^17^. The ANI between these genomes is 96.67% (Fig. 3b), surpassing the species threshold of 95%^45^, suggesting that these genomes represent different strains of the same *Nitrospira* species, potentially adapted to cold environments. In contrast, La3 is grouped within a clade B subset that includes MAGs from drinking water treatment systems and glacier surface soil^46^, but it did not share high ANI values with any other clade B genomes (≤85%, Fig. 3b). This suggests that La3-bin1 represents a unique so far undetected lineage within clade B.

**Fig. 4.**
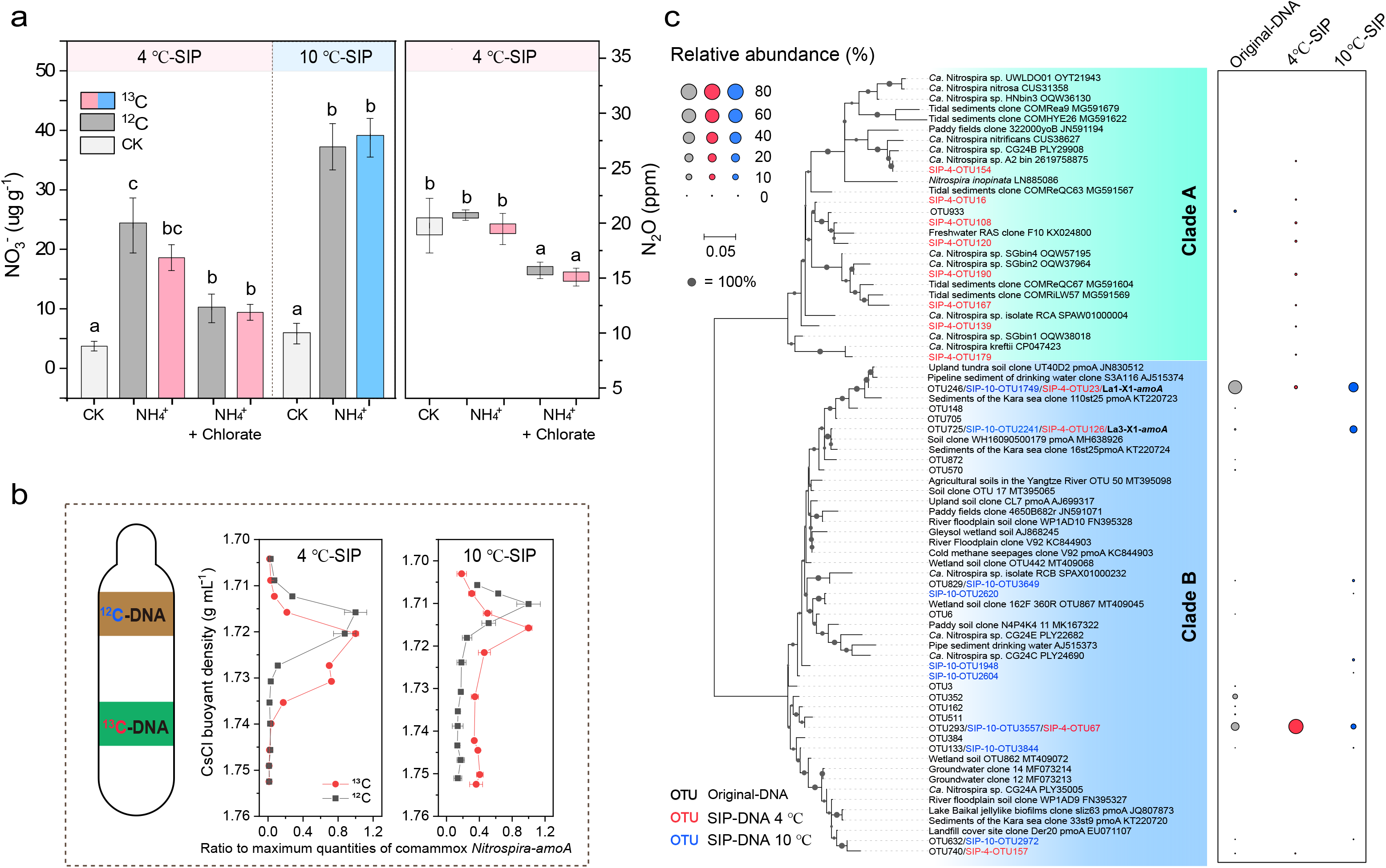
The use of ^13^C-DNA stable isotope probing (SIP) to confirm the nitrification activity of clade B comammox *Nitrospira* in LA1 lake sediment. **(a)** The concentration of produced nitrate (at 4 ℃ and 10 ℃) and N_2_O (at 4 ℃) after 56-day DNA-SIP incubations, using ammonium as substrate and chlorate as specific inhibitor for comammox *Nitrospira*. **(b)** The quantitative distribution and relative abundance of comammox *Nitrospira amoA* genes derived from DNA-SIP in ^13^CO_2_ and ^12^CO_2_-treated microcosms at 4 ℃ and 10 ℃, respectively. Error bars represent standard errors (n = 3). **(c)** Maximum-likelihood phylogenetic trees of retrieved comammox *Nitrospira amoA* gene sequences from the original sample DNA and ^13^C-DNA from 4 ℃ and 10 ℃ DNA-SIP incubation samples. The heatmap on the right displays the relative abundance of specific operational taxonomic units (OTUs) in different DNA samples. The additional DNA-SIP data pertaining to the 10 ℃ incubations of LA2 and LS4, as well as the *nxrB* gene labeling data for LA1, are presented in the Supplementary Fig. 11-13.

### Metabolic potential and surviving strategy of clade B comammox *Nitrospira* in Antarctica

The clade B comammox MAGs *Nitrospira* sp. La1-X1 and *Nitrospira* La3-X1 harbor the complete genetic machinery for NH_3_ and NO_2_^−^ oxidation, the respiratory chain, and the reduced tricarboxylic acid (rTCA) cycle, which is the conserved CO_2_ fixation pathway in *Nitrospira* (Fig. 3c and 3d). These core metabolic features are highly conserved in comammox *Nitrospira* as reported in previous studies^47, 48^. Similar to other *Nitrospira* genomes^46^, La1 and La3 do not encode nitric oxide reductase (NOR), which is crucial for enzymatic N_2_O production^49^. They do carry genes for urea transport and hydrolysis by the urease, indicating the use of urea as an alternative ammonia source, as shown for other comammox *Nitrospira*^9, 10, 50^. In addition, like some *Nitrospira* that have the confirmed ability to use hydrogen and formate as alternative energy sources^51, 52^, the comammox MAGs La1 and La3 contain genes for formate and hydrogen oxidation.

Although the formate dehydrogenases of comammox are similar to their nitrite-oxidizing counterparts, comammox bacteria possess a 3b-type [NiFe] hydrogenase. This hydrogenase is rarely identified in canonical *Nitrospira* and its physiological role remains unclear^48, 53^. While the capacity for formate oxidation is widely distributed in clade B, the 3b-type [NiFe] hydrogenase has been mainly identified in clade A comammox species^48^ (Fig. 3d). However, the presence of this hydrogenase type in a few clade B genomes^46, 48^ including La3 challenges the clade specificity of this feature.

In addition to the canonical F_1_F_0_ H^+^-driven ATPase, La1 and La3 encode a potentially Na^+^-pumping F_1_F_0_ ATPase, previously detected in the haloalkalitolerant nitrite-oxidizing *Ca.* Nitrospira alkalitolerans^54^ and clade A comammox *Ca.* N. kreftii^14^. Similar to these genomes, La3 also possesses a Na^+^-translocating NADH:ubiquinone oxidoreductase (NQR), a feature missing in La1(Fig. 3c). These Na^+^-pumping enzyme complexes might represent an adaption of La3 to saline or haloalkine conditions similar to other *Nitrospira*.

The disaccharide trehalose is one of several solutes known to protect bacteria against cold stress, and may also protect against other harmful environmental conditions, such as osmotic stress^55^. La1 possesses a trehalose-6-phosphate synthase (OtsA) and the corresponding phosphatase (OtsB) to potentially produce the non-reducing disaccharide in two steps from UDP-glucose, as shown for *E. coli*^56^. In contrast, La3 encodes several other trehalose synthesis enzymes in one gene cluster, indicating that this species might convert NDP-glucose (trehalose synthase) and maltose (trehalose synthase/amylase) to trehalose. However, all these trehalose synthesis pathways have also been identified in *Nitrospira* from mesophilic environments and might thus not be a distinguishing feature for cold adaptation. In addition to trehalose synthesis, La1 and La3 possess other features for coping with different stresses. Whereas a gene cluster encoding a nitrile hydratase has been detected in La1 and La3, the genomes lack an amidase to degrade the produced amides further to the corresponding carboxylic acid and ammonia, which could serve as an energy source. An example of a potential amidase is a putative formamidase identified in a clade B MAG obtained from the Rifle aquifer^46^. Other detoxification mechanisms include catalases and Fe orMn superoxide dismutases (SOD) for reactive oxygen defence identified in both clade B genomes, and a periplasmic Cu-Zn SOD present in La3 and many other clade B genomes, but absent in La1 (Fig. 3c and 3d).

An adaptation to oligotrophic conditions (Supplementary Table 1) might be the methionine salvage pathway (MSP) of La1 (Fig. 3c). This pathway recycles the sulphur-containing intermediate 5’-methyl-thioadenosine back to methionine, thus allowing the use of reduced sulphur compounds under sulphur limitation. However, similar to the high-quality genome of the nitrite-oxidizing *Nitrospira lenta* and other clade B genomes, the *mtnE* gene encoding the last step in this pathway is missing in La1. Whether other enzymes might complete the MSP in these *Nitrospira* species remains to be determined.

### Nitrification and N_2_O production activities driven by comammox *Nitrospira*

We have demonstrated that comammox *Nitrospira* potentially serve as crucial drivers of nitrification in coastal East Antarctica, and further investigated their survival strategies from a genomic perspective. Additionally, we confirmed the nitrification activity of comammox *Nitrospira* using DNA-SIP on two lake sediment samples (LA1 and LA2) and one soil sample (LS4). DNA-SIP incubations were conducted at 4°C (for LA1) and 10°C (for LA1, LA2, and LS4) and the production of NO_3_^−^ served as indicator of nitrification activity (Fig. 4a, Supplementary Fig. 11). During the incubations, comammox *Nitrospira* were actively growing, as evidenced by the peak shifts of their DNA the ^13^C-treatment (Fig. 4b, Supplementary Fig. 4). Subsequent sequencing of the *amoA* genes from the labelled DNA revealed clade B comammox *Nitrospira amoA* OTUs, including those of comammox MAGs La1-X1 and La3-X1, in both the 4°C and 10°C incubations (Fig. 4c). Collectively, these data bolster the evidence for the active role of clade B comammox *Nitrospira* in nitrification.

The addition of NH_4_^+^ did increased the nitrate production, however, did not stimulate higher N_2_O production (Fig. 4a). These can be explained by the fact that the major active nitrifiers-comammox *Nitrospira-* are low N_2_O producers^49^. The application of chlorate, a potential comammox-specific inhibitor^57^, slightly reduced the production of NO_3_^−^ and N_2_O (Fig. 4a). However, given that chlorate can also influence other nitrite-oxidizing *Nitrospira* species as well as denitrifiers, it is not feasible to ascribe the observed reductions in NO_3_^−^and N_2_O levels solely to the activity of comammox *Nitrospira*.

Nevertheless, prior research has shown the resilience of clade B comammox *Nitrospira* to freeze-thaw cycles, which underscores the endurance of this particular nitrifier group in environments with low and even subfreezing temperatures^58^. Raising the temperature (10°C) stimulated the activity of AOA (in LA2) and AOB (in LA2 and LS4) (Supplementary Fig.4, Supplementary Results and Discussion), indicating that these nitrifiers are more competitive under elevated temperature conditions, an important finding to predict ecosystem changes in response to global warming.

### Implications and outlook

In this study, we elucidated the unique microbial N-cycle in pristine and oligotrophic coastal Antarctic soil and lake sediments, identifying the microbial nitrification process as the primary pathway for NO_3_^−^ production. Our metagenomic and quantitative functional gene-targeted analyses both revealed clade B comammox *Nitrospira* to be key nitrification drivers in these environments. This finding not only expands our understanding of microbial diversity but also underscores the pivotal role of specific microbial groups in biogeochemical cycling in extreme environments. Our results also revealed fascinating patterns of niche differentiation between clade A and B comammox *Nitrospira* and canonical nitrifiers. The genomic data strongly suggest that clade B species might have superior affinity for their substrate ammonia, and are potentially best adapted to survive and thrive in cold and oligotrophic environments. This niche differentiation could have significant implications for N-cycling in cold environments and it will be important to determine how the nitrifier and overall microbial communities will react to global warming.

In addition, we have successfully obtained in total 1968 MAGs, significantly enriching the existing microbial genomic data for Antarctica. These MAGs provide invaluable insights into the microbial diversity and their metabolic capabilities in the extreme and unique environment of coastal East Antarctica. The availability of these MAGs will undoubtedly facilitate further research into the intricate relationships between microbial communities, biogeochemical cycles, and climate change in polar regions and beyond.

This study underscores the importance of understanding the unique microbial N-cycle in coastal East Antarctica. We have uncovered clade B comammox *Nitrospira* to be a novel nitrification driver in low-temperature environments, and investigated their survival strategy and potential impact on climate change through the production of greenhouse gas N_2_O ^49, 59^. It is crucial to further investigate how comammox *Nitrospira* evolve and survive in Antarctic ecosystems, meeting one of the six priorities for Antarctic science^60^. Recognizing the climatological significance of N-cycling and biogeochemical processes, future research should continue to monitor these microbes, as they could hold the key to understanding the broader implications of microbial activity on our planet’s climate.

## Methods

### Sample collection and treatment

Samples were collected in Larsemann Hills (LH), the second-largest ice-free land in East Antarctica with an area of ∼50 km^2^. LH has a cold and dry continental climate, with an annual mean temperature of ∼-10 °C and the temperatures occasionally above 0 °C in summer^61^, leading to this region typically free of snow cover in summer. As a result of seasonal snow cover and the glacier melting, a number of landlocked lakes are developed in this region. Surface sediment (the upper ∼5 cm) and surface water samples were collected from six lakes (LA1-LA6) in LHs in February 2020 (Fig. 1a). The surface sediment samples were procured using a stainless-steel spade on the shore of lakes, with typical water depths of ∼150-200 cm. Approximately 1L of surface water near the lake shore was also sampled with clean polyethylene (PE) bottles, a portion of which was used to measure temperature, pH, salinity, and conductivity utilizing a multi-probe water quality meter (YSI Professional Plus series; Supplementary Table 1). The remainder of the water samples was filtered using 0.22-μm polytetrafluoroethene (PTFE) filters for chemical analyses. Additionally, a surface soil sample (the top ∼5 cm) was collected near each lake, ∼100 m from the lake shore, using a clean stainless spatula. At each sampling site, larger gravels were removed firstly, and five soil sub-samples (four corners and the center of a square) were collected at a distance of 5–10 m and then mixed to obtain a representative sample (LS1-LS6). For a comprehensive understanding of N cycling processes in the ice-free areas in East Antarctica, two surface soil samples (SVL1 and SVL2) were also collected in February 2022 at Inexpressible Island, South Victoria Land where the climate is comparable to that of LH, following the same sampling protocols. All of the sediment and soil samples were stored in sealed PE bags, and all samples were transported to the laboratory at temperatures of ∼-20 °C for subsequent analysis.

### Physiochemical analysis

In the laboratory, approximately 50 g of the sediment and soil samples were freeze-dried in 50-mL clean centrifuge tubes (ALPHA 1-4/LD, Martin Christ Inc.). After drying, the samples were homogenized using an agate mortar and pestle, and subsequently passed through a 1 mm sieve for further chemical analysis. For determining total organic carbon (TOC) content, approximately 5 g of samples were digested with 10% HCl (*v*/*v*) to remove carbonate. Then, TOC was measured with an automatic element analyser (Elementar, VARIO EL III), with acetanilide used as the external standard. The detection limit (DL) of the TOC was estimated to be approximately 0.005%. All samples were measured in triplicate, yielding a relative standard deviation (1σ) of <10% for each sample. For chemical ion analysis, approximately 5 g of freeze-dried samples were placed in sterile 50-mL centrifuge tubes and suspended in 25 mL Milli-Q water (18.2 MΩ). The solution was then ultrasonicated for 40 min. The supernatant was first centrifuged for 15 min at 3000g, and then filtered through 0.22-μm PTFE filters for nutrient determination. Nutrient concentrations (NH_4_^+^, NO_3_^−^, and PO_4_^3−^) in the sediment, soil extracts, and lake water samples were determined using an Aquion RFIC ion chromatograph (IC, Thermo Scientific, USA), equipped with the analytical columns CS12A (2×250 mm), AS11-HC (2×250 mm), methanesulfonic acid (MSA), and potassium hydroxide (KOH) as eluents for cations and anions, respectively. In addition, the concentrations of SiO_3_^2−^ were determined using an automated QuAAtro™ nutrient analyser (Seal Analytical Ltd., UK). During sample analysis, replicate determinations (*n* = 5) were performed, and 1σ for all species was <5%.

### Isotopic analysis of nitrate

The isotopic composition of NO_3_^−^ was determined using the bacterial denitrifier method at the Environmental Stable Isotope Laboratory of East China Normal University (ECNU-ESIL). Briefly, the denitrifying bacterium *Pseudomonas aureofaciens*, which lacks the N_2_O reductase enzyme, quantitatively transforms NO_3_^−^ into gaseous N_2_O^62, 63^. The δ^15^N and δ^18^O of the generated N_2_O were measured in duplicates using isotope-ratio mass spectrometry (IRMS, Thermo Scientific Delta V). The Δ^17^O of NO_3_^−^ was separately analyzed through the thermal decomposition of N_2_O into to N_2_ and O_2_^64^, followed by measurements at m/z 32, 33, and 34 on the IRMS. The pooled standard deviation (1σ_p_) was employed to ascertain the measurement precision of the overall denitrifier method^65, 66^. The 1σ_p_ of all duplicate samples executed in at least two different batches was 0.6‰ for δ^15^N (n=10), 0.3‰ for δ^18^O (n=10), and 0.8‰ for Δ^17^O (n=8). However, due to the limited amounts of NO_3_^−^ in the samples, only 3 sediment and 5 surface soil samples were analyzed for Δ^17^O of NO_3_^−^.

In addition, lake water stable isotopes (δ^18^O and δ^2^H) were analyzed using laser absorption spectrometry (TIWA-45EP, Los Gatos Research, Inc.). To ensure quality control, replicate analyses (n=5) were performed, yielding relative standard deviations of 0.05‰ and 0.2‰ for δ^18^O and δ^2^H, respectively.

### DNA extraction, PCR, qPCR, sequencing and phylogenetic analysis

Total DNA was extracted from 0.5 g sediment/soil samples using the Fast DNA SPIN kit (MP Biomedicals, Santa Ana, CA) according to the manufacturer’s protocols. The final DNA quality and quantity were determined using Quant-iT PicoGreen dsDNA Assay Kit (ThermoFisher Scientific, China). A detailed description of the Quantitative PCR (qPCR) analysis for functional nitrogen cycling genes was provided in the Supplementary Methods section. It also elaborates on the procedures for PCR and high-throughput sequencing, as well as the subsequent phylogenetic analysis of nitrification genes.

### Metagenomic sequencing

The total DNA from the original sediment and soil samples was sequenced on the Illumina HiSeq Xten platform using a 150-bp paired-end library at Beijing Novogene Biotech Co., Ltd. (Beijing, China). The NEXTFLEX Rapid DNA-Seq Library Prep Kit 2.0 (Bioo Scientific, Austin, TX, USA) was used for DNA library preparation with an insert size of ∼300 bp according to the manufacturer’s recommendations. DNA was sheared using a Covaris S220 Focused Ultrasonicator to create 150 bp fragments. Subsequently, metagenomic sequencing was performed on a cBot Cluster Generation System according to the manufacturer’s standard protocols. we acquired a data range between 20-40 gigabases sequencing data for each individual sample.

### Assembly and binning of metagenomes

Raw reads were processed using fastp v0.19.7^67^ for adapter trimming, quality filtering, and per-read quality trimming. The seven quality-controlled metagenomes were individually assembled and co-assembled using MEGAHIT v1.1.3^68^ with default parameters (k-mers: 21,29,39,59,79,99,119,141). Each of the fourteen assemblies was initially binned using the binning module (– metabat2 –maxbin2 –concoct; –metabat2 for co-assembly) in the metaWRAP pipeline^69^ v1.3.2, and were consolidated using DAS Tool v1.1.2^70^ with default parameters. After dereplication check using dRep v3.0.0^71^ (-comp 50 -con 10) and completeness and contamination evaluation using CheckM v1.1.3^72^, 1968 MAGs were obtained, and the taxonomy of each MAG was assigned using GTDB-Tk v1.5.0^73^ with the Genome Taxonomy Database (Release 06-RS202). In the process of annotating metabolic functions, the genomes that were extracted were examined using DIAMOND^74^ against 52 tailor-made protein databases, which consisted of representative metabolic marker genes. To confirm the existence of crucial metabolic genes in the MAGs, phylogenetic trees were generated using the maximum-likelihood method. The dereplicated MAGs had their percentage relative abundances determined by aligning each sample’s clean paired-end reads to the MAGs utilizing CoverM (version 0.6.1, available at https://github.com/wwood/CoverM) in genome mode, applying the default configurations. Moreover, the trimmed reads were incorporated into CLC Genomics Workbench version 20.0 (CLCBio, Qiagen, Germany), and the de novo assembly algorithm of CLC was employed to search for *amoA* and *nxrB* sequences. Contigs that contained either *amoA* or *nxrB* genes were selected for phylogenetic analysis, in conjunction with amplicon sequences.

### Phylogenomic analysis and genome annotation

All available genomes and MAGs classified by the GTDB-Tk database R202^75^ as *Nitrospiracea*, with estimated genome completeness ≥70% and contamination ≤10%, were downloaded from NCBI. Dereplication was performed using the drep v2.4.2^71^ dereplicate workflow with cut-offs for estimated genome completion ≥70% and contamination ≤10%, but otherwise default settings. In addition to these 95 *Nitrospiracea* genomes, seven *Nitrospira* lineage IV and two clade B genomes were included in the phylogenetic analysis using the UBCG pipeline for the extraction and concatenated alignment of 91 single-copy core genes^76^. In addition, two *Leptospirillum* genomes (GCF_000284315.1, GCF_000299235.1) were included as the outgroup. A maximum-likelihood phylogenetic tree was constructed using IQ-TREE v1.6.12^77^ with 1000 ultrafast bootstrap replications and the GTR+F+I+G4 model identified by the implemented Modelfinder^78^. ANI analysis of all clade B genomes was performed using the OrthoANI^79^. Gene calling and automatic genome annotation were performed using the MicroScope platform, and annotations of selected features were manually checked and refined. For analyzing the distribution patterns of selected key features, 17 comammox clade B genomes were annotated using prokka v.1.14.6^80^ with the “--gcode 11” and “--metagenome” options to obtain all protein sequences for generating a BLAST database. The distribution of selected key proteins within clade B was analyzed by conducting a BLASTp search against this database with default settings, except for an e-value cutoff of 1e-6. Only hits with an identity ≥35% (pident) and a query coverage ≥80% (qcovs) were reported as present. Phylogenomic and ANI analyses were carried out on the retrieved AOA MAGs. The process followed was as previously described, using representative genomes (which include Group I.1a and Group I.1b AOA) and Group I.1b *Nitrosocosmicus* AOA genomes as references, respectively.

### DNA-stable isotope probing (SIP) microcosm incubations, gradient fractionation, quantification, and sequencing

Lake sediments (LA1 and LA2), and the soil sample (LS4) were selected for DNA-SIP microcosm incubation experiments (Supplementary Fig. 10) to mimic a low-temperature oligotrophic environment under in situ flooding conditions. Microcosms were constructed in 120 mL serum bottles containing 20 g sediments at 60% of maximum water-holding capacity and were incubated for 56 days at 10 °C in the dark. For each sample, two different treatments were established in triplicate microcosms. The ^13^CO_2_ microcosms were amended with 5% (*v/v*) ^13^CO_2_ (99 atom%; Sigma-Aldrich Co., St. Louis, MO, USA) plus approximately 5 µg ^15^N-NH_4_Cl-N g^−1^ dry weight soil/sediment (d.w.s.), while the ^12^CO_2_ control treatments received 5% (*v/v*) ^12^CO_2_ plus approximately 5 µg ^14^N-NH_4_Cl-N g^−1^ d.w.s.. The water content was restored weekly over an 8-week incubation period by opening the bottles. The sediment/soil samples were destructively sampled after 56 days of incubation and transferred immediately to –80°C for subsequent molecular analysis. The remaining ∼2 g of sediments were used for end-point quantification of NH_4_^+^, NO_2_^−^, and NO_3_^−^ concentrations. The LA1 sediment sample was additionally chosen for an incubation at 4 °C, adhering to the same procedure previously outlined. In addition to the ammonium substrate, 50 μM chlorate, a specific comammox inhibitor^57^, was introduced to study the activity of comammox *Nitrospira*.

Moreover, the production of nitrous oxide (N_2_O)^57^ was actively monitored during these 4°C DNA-SIP incubations. The fractionation of DNA post DNA-SIP incubations, along with the subsequent quantification analysis of functional nitrification groups (detailed in the Supplementary Methods).

### Data availability

All sequences of AOA-, AOB-, and comammox *Nitrospira-amoA* and *Nitrospira-nxrB* obtained in this study were deposited in GenBank, with accession numbers MZ956347-MZ956585. All amplicon sequencing data, unprocessed metagenomes, metagenomic assemblies, and metagenome-assembled genomes (MAGs) have been submitted to the Sequence Read Archive of the National Center for Biotechnology Information (NCBI), under the BioProject accession No. PRJNA855145. The sequences and associated annotations of the five AOA and seven *Nitrospira* MAGs are publicly available at MicroScope (https://mage.genoscope.cns.fr/microscope).

## Acknowledgements

This work was supported by the National Natural Science Foundation of China (NSFC) (Nos. 42371064, 42030411, 42276243, 42030411, and 41807465), and the Program of Shanghai Academic/Technology Research Leader (Grant No. 20XD1421600), and by the Dutch Research Council (NWO; Talent Programme grants VI.Veni.192.086 and 016.Vidi.189.050). The LABGeM (CEA/Genoscope & CNRS UMR8030), the France Génomique and French Bioinformatics Institute national infrastructures (funded as part of Investissement d’Avenir program managed by Agence Nationale pour la Recherche, contracts ANR-10-INBS-09 and ANR-11-INBS-0013) are acknowledged for support within the MicroScope annotation platform.

## Author contributions

P.H., G.S. and M.L. conceived the study. P.H. and X.T. conducted the lab work. P.H. and H.K. performed the comparative genomic analyses. X.D. and P.H. analyzed the metagenomic data. D.W. and Z.L. collected the samples. Q.Z. did the physiochemical analyses. L.H. and S.L. helped with the data interpretation. P.H. and G.S. wrote the manuscript, with contributions from all co-authors.

