## Supplementary Information for "Unveiling unique microbial nitrogen cycling and novel nitrification drivers in coastal Antarctica"

### Supplementary Results and Discussion

**Abundant and diverse comammox *Nitrospira*.** We conducted an analysis of the abundance and community composition of ammonia-oxidizing archaea (AOA), ammonia-oxidizing bacteria (AOB), and comammox *Nitrospira* in the sediments of six distinct lakes (LA1 to LA6) and a proximal soil sampling location (LS4) (Supplementary Table 1). This analysis centered on the *amoA* gene, which encodes the alpha subunit of ammonia monooxygenase and serves as a key phylogenetic marker for ammonia oxidizers. For comammox *Nitrospira*, the presence of clades A and B was evaluated using clade-specific *amoA*-targeting primers[1]. AmoA gene quantification revealed that clade B comammox *Nitrospira* represented over 90% of the total ammonia-oxidizing population, indicating their dominance within the ammonia-oxidizing community and surpassing clade A comammox *Nitrospira,* AOA, and AOB (Supplementary Fig. 3).

Additionally, we assessed the overall abundance and diversity of *Nitrospira*, which includes both strict nitrite-oxidizing species and comammox *Nitrospira*, by targeting the *nxrB* gene. This gene encodes the beta subunit of the nitrite oxidoreductase, which is crucial for nitrite oxidation. The *nxrB* gene abundance was found to be on par with that of comammox *Nitrospira amoA* (Supplementary Fig. 3), implying that the majority of *Nitrospira* present in these lake sediments are likely capable of performing complete nitrification.

The AOA *amoA* OTUs we detected were predominantly part of the Group I.1b

*Nitrosocosmicus*-cluster (Supplementary Fig. 4), whereas the AOB *amoA* OTUs were chiefly classified within the genus *Nitrosospira* (Supplementary Fig. 7). These findings are in alignment with prior studies [2-4]. Across all lake sediments and the soil sample LS4, we observed a consistent community structure for AOA, AOB, and comammox *Nitrospira* (Supplementary Fig. 8).

We observed a significantly greater diversity of comammox *Nitrospira* compared to that of AOA and AOB, as evidenced by the number of *amoA* operational taxonomic units (OTUs) identified (Supplementary Table 4). The diversity of *Nitrospira*, as determined by *nxrB* gene sequences, was found to be on par with that of comammox *Nitrospira*. This finding reinforces the notion of comammox *Nitrospira*'s predominance over traditional nitrifiers, including the strict nitrite-oxidizing *Nitrospira* species. The majority of comammox *Nitrospira amoA* OTUs were associated with clade B (Supplementary Fig. 8). Specifically, OTU246, which was most frequently detected in lake LA1, and OTU725, prevalent in other samples, emerged as the two most abundant comammox *amoA* OTUs. Similarly, two distinct *Nitrospira nxrB* OTUs were observed, with OTU70 being predominantly present in sample LA1, and OTU179 being prevalent in the rest of the samples (Supplementary Fig. 9). Intriguingly, the *amoA* sequences obtained from the two metagenome-assembled genomes (MAGs) of comammox *Nitrospira*, designated La1 and La3, were an exact match to these two dominant *amoA* OTUs (Supplementary Fig. 8). This suggests that these particular comammox *Nitrospira* strains are the leading nitrifiers within these aquatic ecosystems. Similarly, the *nxrB* sequence from the MAG La1 and a *nxrB*-bearing contig from the LA3 lake assembly were identical to the two most abundant *nxrB* OTUs, further substantiating their significance in the nitrification process within these environments.

Regarding the total *Nitrospira* population, members of lineage II, which includes both comammox and nitrite-oxidizing species, were preeminent in all lake sediment samples. In stark contrast, *Nitrospira* from lineage IV, typically encountered in marine or coastal settings, constituted a significant majority (>75%) of the total *Nitrospira* population in the soil sample LS4 (Supplementary Fig. 9). The prevalence of lineage IV in this context suggests that the surface soil experiences greater influence from sea salt aerosols, derived from the neighboring Antarctic Ocean (Fig. 1a), than does the lake sediment. Moreover, this dominance implies that lineage IV *Nitrospira* may be more adept at thriving in terrestrial habitats[5] than previously recognized.

**Active nitrifiers revealed by $^{13}C$-DNA-SIP.** Following microcosm incubations with both unlabelled ($^{12}CO_2$ and $^{14}NH_4Cl$) and labelled ($^{13}CO_2$ and $^{15}NH_4Cl$) substrates (Supplementary Fig. 10), conducted at temperatures of 4°C and 10°C, we noted $NO_3^-$ accumulation in all samples from LA1 at both temperatures, in LA2 at 10°C, and in LS4 at 10°C (Figure 4a and Supplementary Fig. 11a). This accumulation is indicative of active nitrification processes. Comammox *Nitrospira* were present in all samples, and AOB were identified in some, while AOA were absent (Supplementary Fig. 11a). Specifically, in the LA1 samples, comammox *Nitrospira* were the sole nitrifiers detected in the incubations. In contrast, for LA2 and LS4 samples, both comammox

*Nitrospira* and AOB were identified (Supplementary Fig. 11b). In alignment with these observations based on *amoA* gene presence, the *nxrB* gene of *Nitrospira* was found in all the samples tested (Supplementary Fig. 11b), confirming the widespread distribution of nitrifying *Nitrospira* across the different environments sampled.

Based on quantification of the functional genes, peak shifts in $^{13}$C-DNA compared to $^{12}$C-DNA were observed in specific *amoA* and *nxrB*-based comammox and total *Nitrospira* abundances, respectively, for all tested samples (Fig. 4a, Supplementary Fig. 12). This finding confirmed the activity of *Nitrospira* and their importance in nitrification and thus $NO_3^-$ production in Antarctic Lake sediments. Notably, although AOB were detected in both LA2 and LS4 based on AOB-specific *amoA* PCR, their activity was only confirmed for LS4 (Supplementary Fig. 4a, Supplementary Fig. 12), further illustrating and supporting the different nitrifier compositions and contributions between surface soil and lake sediments in Antarctica. This difference also aligns with a recent study in Antarctic desert soils where Thermoproteota (AOA) and *Nitrospirota* were the most abundant ammonia and nitrite oxidizers, respectively[6]. According to relative *amoA* abundances, the dominant comammox OTUs were in accordance with those identified in the original samples (Supplementary Fig. 8), proving that the most abundant comammox *Nitrospira* drive the nitrification activity in soils/lake sediments tested. For LA1 and LA2, the *nxrB* data showed similar concordance between the original and $^{13}$C-labelled DNA, further supporting the dominance of comammox *Nitrospira* in ammonia and nitrite oxidation (Supplementary Fig. 13). Notably, lineage IV *Nitrospira* again dominated over lineage

II in the $^{13}$C-DNA from the LS4 soil incubation. At this site, the additional ammonia oxidation activity by AOB might provide more nitrite than leaked by comammox *Nitrospira* (Supplementary Fig. 13), which may stimulate the activity and growth of nitrite-oxidizing lineage IV *Nitrospira*.

### Supplementary Methods

**Quantitative PCR (qPCR) analysis.** The abundances of functional nitrogen cycling genes *amoA* genes, including nitrogen fixation (*nifH*), nitrification (*amoA, hao*, and *nxrB*), denitrification (*napA*, *narG*, *nirS/K*, *norB*, and *nosZ*), dissimilatory nitrate reduction to ammonium (DNRA; *nrfA*), assimilatory nitrogen reduction (ANRA; *nasA* and *nirA*), and anaerobic ammonium oxidation (Anammox; *hzo*), were quantified by qPCR on an ABI 7500 Real-Time PCR System (Applied Biosystems, Canada) in 21 µL reaction mixtures. The mixtures contain the following components: 10 µL of Hieff® qPCR SYBR Green Master Mix with Low Rox Plus (Yeasan, China), 0.4 µL of each primer (50 µM), 1 µL of template DNA and 9.2 µL of $ddH_2O$. The primer sets and amplification conditions are listed in Supplementary Table 2. Standards for quantifying gene abundance were generated from the respective purified PCR products using the QIAquick PCR Purification Kit (Qiagen, Germany) following the manufacturer's protocol. The concentration of the purified PCR products was estimated using Quant-iT PicoGreen dsDNA Assay Kit (ThermoFisher Scientific, China), and converted to gene copy numbers.

**PCR, high-throughput sequencing and phylogenetic analysis.** The *amoA* genes of

AOA, AOB, and comammox *Nitrospira* were amplified using the primer sets CamoA-19F/616R (AOA)[7], amoA1F/2R (AOB)[8], comaA-244f/659r (comammox clade A)[1], comaB-244f/659r (comammox clade B)[1], and Ntsp-amoA-162F/359R (general comammox)[9], respectively. *Nitrospira nxrB* genes were amplified using the primer set nxrB169F/638R[10]. Primer sequences and PCR conditions are listed in Supplementary Table 2. PCR was performed in 50 μL volume system containing 25 μL of 5× Green Taq Mix (Vazyme, Nanjing, China), 1 μL of each primer (10 μM), 2 μL of template DNA, and 21 μL of dd$H_2O$. PCR products were purified by 1.5% agarose gel electrophoresis and quantified using the Picogreen quantification kit (Invitrogen, Shanghai, China). After the first round of functional gene PCR screening for nitrifiers, we used barcoded primers for subsequent PCR amplification and high-throughput sequencing. In brief, 10 bp barcodes were added to each forward primer to differentiate the samples. After PCR amplification and product purification, PCR products of the *amoA* genes of AOA, AOB, and comammox *Nitrospira* (with primer set Ntsp-amoA-162F/359R), and *nxrB* genes of *Nitrospira* were paired-end sequenced using an Illumina MiSeq platform (2×300 bp) by Majorbio Bio-Pharm Technology Co. Ltd. (Shanghai, China) according to the manufacturer's protocols.

Raw sequencing reads were checked for quality using FastQC (http://www.bioinformatics.babraham.ac.uk/projects/fastqc/) and trimmed using the Quantitative Insight into Microbial Ecology (QIIME) pipeline[11]. The paired-end reads were merged using FLASH[12] and filtered based on their Phred quality scores (Q=20). Chimeric sequences were identified using de novo detection with the flag "--

non_chimeras_rentention"[13]. The remaining high-quality sequences were clustered into operational taxonomic units (OTUs) using UCLUST based on a 95% similarity cutoff.

The length of the AOA *amoA* obtained using primers CamoA-19F/616R was longer than the read length from paired-end sequencing (2×300 bp). We speculated that the deletion of 30 nucleotides in the middle of the AOA *amoA* fragments would not significantly influence phylogenetic analysis, as evidenced by the phylogenetic comparison of representative AOA *amoA* sequences with and without the 30 nucleotides. First, every single read was quality-trimmed using fastp[14] with a sliding window of 50 bp. When the average quality score within the window dropped below 30, all subsequent bases were trimmed. Only reads ⩾ 260 bp were retained. Then, the non-overlapped paired-end reads generated from the two opposite ends of the same DNA fragment were directly joined into one sequence and aligned to representative sequences of AOA *amoA* genes from the NCycDB database[15] using hmmalign[16]. In the alignment, the ~30 bp non-hit region at the joint of paired-end reads was trimmed for phylogenetic analysis.

For phylogenetic analyses, the representative sequences of each OTU and reference sequences from each nitrifying guild were aligned using hmmalign[16], and the alignments were trimmed using TrimAL[17] with the "-gappyout" flag. Maximum likelihood trees were constructed using IQ-TREE[18] with the "-MFP -bb 1000" flags for best-fit model selection and 1000 ultrafast bootstraps[19].

**Statistical analysis.** The number of *amoA* and *nxrB* genes for ammonia oxidizers and *Nitrospira*, respectively, were analyzed using Origin 2018 (OriginLab, USA). Different metrics of alpha diversity (observed OTUs, Chao1, Shannon, Simpson, and phylogenetic diversity) were computed using QIIME (version 2)[11]. Statistical correlations between sediment properties and comammox *Nitrospira amoA* gene abundances were analyzed through canonical correspondence analysis (CCA) using the vegan package of R 3.4.1.

**DNA-SIP fractionation.** The fractionation of DNA extracted from the SIP incubations was conducted as described previously[20]. Briefly, total DNA of sediment/soil was extracted from 0.5 g sample material using the FastDNA Spin Kit for Soil (MP Biomedicals, Cleveland, OH, USA) according to the manufacturer's instructions. For each treatment, ~3 μg of the extracted DNA was mixed with the CsCl stock solution with an initial density of 1.725 g $mL^{-1}$ in Tris-EDTA (TE) buffer (pH = 8.0). The isopycnic density centrifugation was performed in 5.1 ml Quick-Seal polyallomer ultracentrifugation tube in a VTi65.2 vertical rotor (Beckman Coulter Inc., Palo Alto, CA, USA). The heavy fractions of $^{13}C$-labelled DNA were resolved from the light fractions by ultracentrifugation at 177,000 × *g* for 44 h at 20°C. The resulting gradients were fractionated into 15 equal volumes (approximately 340 μL each) by displacing the gradient medium with sterile water from the top of the ultracentrifuge tube using a syringe pump (New Era Pump Systems Inc., Farmingdale, NY, USA), with a precisely controlled flow rate of 0.34 mL $min^{-1}$. The buoyant density of each fraction was calculated using an AR200 digital hand-held refractometer (Reichert Inc., Buffalo, NY, USA) by measuring the refractive index of 65 μL aliquots. Subsequently, the fractionated DNA was precipitated, washed twice with 70% ethanol, and dissolved in 30 μL TE buffer as described previously[21]. The abundances of *amoA* and *nxrB* of comammox and all *Nitrospira* were quantified using the primer sets Ntsp-amoA-162F/359R and nxrB169F/638R (Supplementary Table 2), respectively. For the heavy fractions with the highest amount of comammox *Nitrospira amoA* and *Nitrospira nxrB*, the obtained PCR products were sent for high-throughput amplicon sequencing, with subsequent phylogenetic analysis as described above.

**Supplementary Table 1.** Details of the sampling locations and geochemical data of lake sediments and overlying waters, and surface soil samples from Larsemann Hills (LH) in coastal East Antarctica, as well as surface soil samples from South Victoria Land (SVL). n.a., not available.

| Location | Larsemann Hills | | | | | | South Victoria Land | |
|---|---|---|---|---|---|---|---|---|
| **Lake sediment** | **LA1** | **LA2** | **LA3** | **LA4** | **LA5** | **LA6** | | |
| Longitude (E) | 76°19' | 76°22' | 76°23' | 76°23' | 76°21' | 76°20' | | |
| Latitude (S) | 69°24' | 69°24' | 69°23' | 69°22' | 69°24' | 69°24' | | |
| TOC/% | 0.07 | 0.03 | 0.17 | 0.07 | 0.03 | 0.04 | | |
| $NH_4^+$ /μg $g^{-1}$ | 0.65 | 1.84 | 5.96 | 2.64 | 2.29 | 1.35 | | |
| $NO_3^-$ /μg $g^{-1}$ | 0.45 | 0.16 | 0.47 | 0.46 | 4.9 | 0.21 | | |
| $SiO_3^{2-}$ /μg $g^{-1}$ | 1.62 | 1.07 | 0.60 | 1.70 | 1.74 | 0.99 | | |
| $PO_4^{3-}$ /μg $g^{-1}$ | 1.06 | 0.79 | 2.18 | 0.07 | 1.41 | 0.11 | | |
| $\delta^{15}N$-$NO_3^-$/‰ | -3.4 | 0.28 | -0.39 | -3.17 | -5.9 | 2.4 | | |
| $\delta^{18}O$-$NO_3^-$/‰ | 7.87 | 7.08 | 8.5 | -5.4 | -16.04 | 25.6 | | |
| $\Delta^{17}O$-$NO_3^-$/‰ | 1.30 | 2.01 | n.a. | n.a. | 0.61 | n.a. | | |
| **Lake water** | **LA1** | **LA2** | **LA3** | **LA4** | **LA5** | **LA6** | | |
| Temperature/$^o$C | 1.5 | 1.8 | 2 | 1.5 | 1.2 | 0.1 | | |
| pH | 6.32 | 6.95 | 7.63 | 7.66 | 6.96 | 7.1 | | |
| Salinity/‰ | 0.06 | 0.09 | 0.56 | 0.52 | 0.08 | 0.5 | | |
| Conductivity/mS $cm^{-1}$ | 0.13 | 0.18 | 1.13 | 1.07 | 0.17 | 1.03 | | |
| $NH_4^+$ /μmol $L^{-1}$ | 2.43 | 2.41 | 3.56 | 2.93 | 1.52 | 1.86 | | |
| $NO_3^-$ /μmol $L^{-1}$ | 1.17 | 0.89 | 0.92 | 1.01 | 0.88 | 0.79 | | |
| $SiO_3^{2-}$ /μmol $L^{-1}$ | 0.02 | 2.30 | 11.51 | 12.15 | 2.97 | 6.96 | | |
| $PO_4^{3-}$ /μmol $L^{-1}$ | 0.13 | 0.06 | 0.13 | 0.12 | 0.01 | 0.03 | | |
| $\delta^{15}N$-$NO_3^-$/‰ | 4.87 | 9.62 | 1.40 | 6.38 | 2.78 | 1.70 | | |
| $\delta^{18}O$-$NO_3^-$/‰ | 22.27 | 5.58 | 11.98 | 3.75 | 17.51 | 28.96 | | |
| $\delta^{18}O$-$H_2O$/‰ | -14.62 | -14.35 | -10.57 | -11.8 | -12.23 | -12.86 | | |
| $\delta^{2}H$-$H_2O$/‰ | -104.88 | -101.51 | -85.21 | -92.72 | -93.4 | -96.67 | | |
| **Soil** | **LS1** | **LS2** | **LS3** | **LS4** | **LS5** | **LS6** | **SVL1** | **SVL2** |
| Longitude (E) | 76°19' | 76°22' | 76°23' | 76°23' | 76°21' | 76°20' | 163°42.54′ | 163°42.61′ |
| Latitude (S) | 69°24' | 69°24' | 69°23' | 69°22' | 69°24' | 69°24' | 74°56.12′ | 74°56.01′ |
| TOC/% | 0.03 | 0.02 | 0.02 | 0.10 | 0.11 | 0.09 | 0.13 | 0.09 |
| $NH_4^+$ /μg $g^{-1}$ | 4.15 | 3.61 | 4.86 | 0.88 | 3.52 | 1.56 | 0.41 | 0.83 |
| $NO_3^-$ /μg $g^{-1}$ | 2.92 | 0.67 | 1.01 | 0.34 | 0.25 | 0.26 | 2.82 | 3.62 |
| $SiO_3^{2-}$ /μg $g^{-1}$ | 6.87 | 2.42 | 0.55 | 2.44 | 3.83 | 3.23 | 3.26 | 10.27 |
| $PO_4^{3-}$ /μg $g^{-1}$ | 1.09 | 1.59 | 0.26 | 1.69 | 8.61 | 0.48 | 0.44 | 2.83 |
| $\delta^{15}N$-$NO_3^-$/‰ | 0.28 | -3.36 | -4.94 | 7.77 | -0.31 | 3.86 | -4.52 | -1.85 |
| $\delta^{18}O$-$NO_3^-$/‰ | 6.20 | -5.14 | 4.48 | -7.47 | 4.94 | 13.91 | 1.24 | 1.76 |
| $\Delta^{17}O$-$NO_3^-$/‰ | n.a. | 0.00 | 0.48 | 0.40 | n.a. | n.a. | 0.98 | 0.72 |

**Supplementary Table 2.** Characteristics of primer sets, including the thermal PCR and qPCR profile, used for the amplification and analysis of functional genes involved in microbial nitrogen cycling processes. These processes include nitrogen fixation (*nifH*), nitrification (*amoA, hao*, and *nxrB*), denitrification (*napA*, *narG*, *nirS/K*, *norB*, and *nosZ*), dissimilatory nitrate reduction to ammonium (DNRA; *nrfA*), assimilatory nitrogen reduction (ANR; *nasA* and *nirA*), and anaerobic ammonium oxidation (anammox; *hzo* and *hzs*).

| | **Target gene** | **Primer name** | **Forward primer (5'-3')** [a] | **Thermal profile for PCR** | **Thermal profile for qPCR** | **Length (bp)** | **Reference** |
|---|---|---|---|---|---|---|---|
| **Nitrification** | **AOA-*amoA*** | CamoA-19F<br>CamoA-616R | ATGGTCTGGYTWAGACG<br>GCCATCCABCKRTANGTCCA | 95 °C for 5 min, followed by 35 cycles of 15 s at 95 °C, 30 s at 53 °C and 45 s at 72 °C. | 50 °C for 2 min and 95 °C for 10 min, followed by 40 cycles of 15 s at 95 °C, 30 s at 53 °C and 40 s at 72 °C. | 629 | 22 |
| | **AOB-*amoA*** | amoA1F<br>amoA2R | GGGGTTTCTACTGGTGGT<br>CCCCTCTGCAAAGCCTTCTTC | 95 °C for 5min, followed by 35 cycles of 15 s at 95 °C, 30 s at 54 °C and 45 s at 72 °C. | 50 °C for 2 min and 95 °C for 10 min, followed by 40 cycles of 15 s at 95 °C, 30 s at 54 °C and 40 s at 72 °C. | 491 | 8 |
| | **Comammox-*amoA*** | Ntsp-amoA-162F<br>Ntsp-amoA-35R | GGATTTCTGGNTSGATTGGA<br>WAGTTNGACCACCASTACCA | 95 °C for 1 min, followed by 35 cycles of 10 s at 95 °C, 40 s at 48 °C and 45 s at 72 °C. | 50 °C for 2 min and 95 °C for 10 min, followed by 40 cycles of 15 s at 95 °C, 30 s at 48 °C and 45 s at 72 °C. | 198 | 9 |
| | **Comammox-clade A *amoA*** | comaA-244f_a<br>comaA-244f_b<br>comaA-244f_c<br>comaA-244f_d<br>comaA-244f_e<br>comaA-244f_f<br>comaA-659r_a<br>comaA-659r_b<br>comaA-659r_c<br>comaA-659r_d<br>comaA-659r_e<br>comaA-659r_f | TACAACTGGGTGAACTA<br>TATAACTGGGTGAACTA<br>TACAATTGGGTGAACTA<br>TACAACTGGGTCAACTA<br>TACAACTGGGTCAATTA<br>TATAACTGGGTCAATTA<br>AGATCATGGTGCTATG<br>AAATCATGGTGCTATG<br>AGATCATGGTGCTGTG<br>AAATCATGGTGCTGTG<br>AGATCATCGTGCTGTG<br>AAATCATCGTGCTGTG | 95 °C for 1 min, followed by 35 cycles of 10 s at 95 °C, 30 s at 53 °C and 45 s at 72 °C. | 50 °C for 2 min and 95 °C for 10 min, followed by 40 cycles of 15 s at 95 °C, 30 s at 53 °C and 45 s at 72 °C. | 415 | 1 |

| | | | | | | | |
|---|---|---|---|---|---|---|---|
| | **Comammox-clade B *amoA*** | comaB-244f_a<br>comaB-244f_b<br>comaB-244f_c<br>comaB-244f_d<br>comaB-244f_e<br>comaB-244f_f<br>comaB-659r_a<br>comaB-659r_b<br>comaB-659r_c<br>comaB-659r_d<br>comaB-659r_e<br>comaB-659r_f | TAYTTCTGGACGTTCTA<br>TAYTTCTGGACATTCTA<br>TACTTCTGGACTTTCTA<br>TAYTTCTGGACGTTTTA<br>TAYTTCTGGACATTTTA<br>TACTTCTGGACCTTCTA<br>ARATCCAGACGGTGTG<br>ARATCCAAACGGTGTG<br>ARATCCAGACAGTGTG<br>ARATCCAAACAGTGTG<br>AGATCCAGACTGTGTG<br>AGATCCAAACAGTGTG | 95 °C for 1 min, followed by 35 cycles of 10 s at 95 °C, 30 s at 53 °C and 45 s at 72 °C. | 50 °C for 2 min and 95 °C for 10 min, followed by 40 cycles of 15 s at 95 °C, 30 s at 53 °C and 45 s at 72 °C. | 415 | |
| | ***hao*** | hao1F<br>hao2R | TGAGCCAGTCCAACGTGCAT<br>GCAACAACCCTGCCT CA | 95 °C for 1 min, followed by 35 cycles of 30 s at 95 °C, 30 s at 58 °C, 1 min at 72 °C. | 50 °C for 2 min and 95 °C for 10 min, followed by 40 cycles of 15 s at 95 °C, 30 s at 58 °C and 1 min at 72 °C. | 219 | 23 |
| | ***Nitrospira-nxrB*** | nxrB169F<br>nxrB638R | TACATGTGGTGGAACA<br>CGGTTCTGGTCRATCA | 95 °C for 5 min, followed by 35 cycles of 40 s at 95 °C, 40 s at 56.2 °C and 90 s at 72 °C. | 50 °C for 2 min and 95 °C for 10 min, followed by 40 cycles of 15 s at 95 °C, 40 s at 56.2 °C and 90 s at 72 °C. | 469 | 24 |
| | ***hzo*** | hzocl1F1<br>hzocl1R2 | TGYAAGACYTGYCAYTGG<br>ACTCCAGATRTGCTGACC | 95 °C for 1 min, followed by 35 cycles of 10 s at 95 °C, 30 s at 50 °C, 30 s at 72 °C. | 50 °C for 2 min and 95 °C for 10 min, followed by 40 cycles of 15 s at 95 °C, 30 s at 50 °C and 30 s at 72 °C. | 994 | 23 |
| **Anammox** | ***hzsA*** | hzsA_526F<br>hzsA_1857R | TAYTTTGAAGGDGACTGG<br>AAABGGYGAATCATARTGGC | 95 °C for 1 min, followed by 35 cycles of 30 s at 95 °C, 45 s at 54°C, 1 min at 72 °C. | 50 °C for 2 min and 95 °C for 10 min, followed by 40 cycles of 15 s at 95 °C, 45 s at 54 °C and 1 min at 72 °C. | 1231 | |
| | ***hzsB*** | hzsB_396F<br>hzsB_742R | ARGGHTGGGGHAGYTGGAAG<br>GTYCCHACRTCATGVGTCTG | 95 °C for 1 min, followed by 35 cycles of 1 min at 95 °C, 1 min at 59 °C, 45 s at 72 °C. | 50 °C for 2 min and 95 °C for 10 min, followed by 40 cycles of 15 s at 95 °C, 1 min at 59 °C and 45 s at 72 °C. | 346 | 25 |
| | ***hzsC*** | hzsC745f<br>hzsC862r | CCRAAGAACTGGYTDCCKGTDTG<br>TAHGGATTNCCRTCRTARTTRTT | 95 °C for 1 min, followed by 35 cycles of 45 s at 95 °C, 45 s at 55 °C, 50 s at 72 °C. | 50 °C for 2 min and 95 °C for 10 min, followed by 40 cycles of 15 s at 95 °C, 45 s at 55 °C and 50 s at 72 °C. | 117 | |
| **Assimilatory nitrate** | ***nasA*** | *nasA*1735 | ATNGTRTGCCAYTGRTC | 95 °C for 1 min, followed by 35 cycles of 1 min at 93 °C, 20 s at | 50 °C for 2 min and 95 °C for 10 min, followed by 40 cycles of 15 s at 95 °C, | 198 | 26 |

| | | | | | | | |
|---|---|---|---|---|---|---|---|
| **reduction** | | *nasA*1933 | CARTGCATNGGNAYRAA | 55.5 °C, 1 min at 72 °C. | 20 s at 55.5 °C and 1 min at 72 °C. | | |
| | ***narB*** | narB430 | AACACIACGCTGTGTATGGC | 95 °C for 1 min, followed by 35 cycles of 1 min at 93 °C, 1 min at 57 °C, 1 min at 72 °C. | 50 °C for 2 min and 95 °C for 10 min, followed by 40 cycles of 1 min at 93 °C, 1 min at 57 °C and 1 min at 72 °C. | 589 | 27 |
| | | narB1019 | GARTTIGCCTGICCGGTCA | | | | |
| | ***nirA*** | nirA136 | TGGTGGGGICTITWCCACCAG | 95 °C for 1 min, followed by 35 cycles of. | 50 °C for 2 min and 95 °C for 10 min, followed by 40 cycles of 15 s at 95 °C, 45 s at 50 °C and 1 min at 72 °C. | 537 | 27 |
| | | nirA673 | CGACRCGRACRTTGAABCC | | | | |
| **DNRA/Denitrification** | ***narG*** | 1960m2f | TAGTGGGCAG GAAAACTG | 95 °C for 1 min, followed by 35 cycles of 15 s at 95 °C, 30 s at 63°C, 30 s at 72 °C. | 50 °C for 2 min and 95 °C for 10 min, followed by 40 cycles of 15 s at 95 °C, 30 s at 63 °C and 30 s at 72 °C. | 90 | 28 |
| | | 2050m2r | CGTAGAAGAAGCTGGTGCTGTT | | | | |
| **DNRA** | ***nrfA*** | nrfAF1 | GCNTGYTGGWSNTGYAA | 95 °C for 1 min, followed by 35 cycles of 30 s at 94 °C, 30 s at 45 °C, 1 min at 72 °C. | 50 °C for 2 min and 95 °C for 10 min, followed by 40 cycles of 30 s at 94 °C, 30 s at 45 °C and 1 min at 72 °C. | 500 | 29 |
| | | nrfAR1 | TWNGGCATRTGRCARTC | | | | |
| | ***nirS*** | cd3aF | GTSAACGTSAAGGARACSGG | 95 °C for 1 min, followed by 35 cycles of 30 s at 95 °C, 40 s at 58 °C, 1 min at 72 °C. | 50 °C for 2 min and 95 °C for 10 min, followed by 40 cycles of 15 s at 95 °C, 40 s at 58 °C and 45 s at 72 °C. | 525 | 30 |
| | | R3cd | GASTTCGGRTGSGTCTTGA | | | | |
| | ***nirK*** | nirK876 | ATYGGCGGVAYGGCGA | 95 °C for 1 min, followed by 35 cycles of 30 s at 95 °C, 15 s at 57 °C, 1 min at 72 °C. | 50 °C for 2 min and 95 °C for 10 min, followed by 40 cycles of 15 s at 95 °C, 15 s at 57 °C and 45 s at 72 °C. | 164 | 31 |
| **Denitrification** | | nirK1040 | GCCTCGATCAGRTTRTGGTT | | | | |
| | ***nosZ*** | nosZ2F | CGCRACGGCAASAAGGTSMSSGT | 95 °C for 1 min, followed by 35 cycles of 30 s at 95 °C, 40 s at 58 °C, 1 min at 72 °C. | 50 °C for 2 min and 95 °C for 10 min, followed by 40 cycles of 15 s at 95 °C, 40 s at 58 °C and 45 s at 72 °C. | 267 | 32 |
| | | nosZ2R | CAKRTGCAKSGCRTGGCAGAA | | | | |
| | ***norB*** | norB1f | CGNGARTTYCTSGARCARCC | 95 °C for 1 min, followed by 35 cycles of 1 min at 93 °C, 1 min at 55 °C, 1 min at 72 °C. | 50 °C for 2 min and 95 °C for 10 min, followed by 40 cycles of 15 s at 95 °C, 1 min at 55 °C and 1 min at 72 °C. | 669 | 28 |
| | | norB8r | CRTADGCVCCRWAGAAVGC | | | | |
| **Nitrogen fixation** | ***nifH*** | nifHF | AAAGGYGGWATCGGYAARTCCACCAC | 95 °C for 1 min, followed by 35 cycles of 10 s at 95 °C, 30 s at 55 °C, 30 s at 72 °C. | 50 °C for 2 min and 95 °C for 10 min, followed by 40 cycles of 15 s at 95 °C, 30 s at 55 °C and 30 s at 72 °C. | 342 | 33 |
| | | nifHR | TTGTTSGCSGCRTACATSGCCATCAT | | | | |

| | | | | | | | |
|---|---|---|---|---|---|---|---|
| **Aerobic denitrification** | ***napA*** | v66 | TAYTTYYTNHSNAARATHATGTAYGG | 95 °C for 1 min, followed by 35 cycles of 1 min at 94 °C, 1 min at 50 °C, 2 min at 72 °C. | 50 °C for 2 min and 95 °C for 10 min, followed by 40 cycles of 15 s at 95 °C, 1 min at 50 °C and 2 min at 72 °C. | 707 | 34 |
| | | v67 | DATNGGRTGCATYTCNGCCATRTT | | | | |

**Supplementary Table 3.** Features of the retrieved MAGs including ammonia-oxidizing archaea (AOA), complete ammonia oxidizers (comammox), and nitrite-oxidizing bacteria (NOB) of the *Nitrospira* genus, as reconstructed from Antarctic metagenomes. The table also includes data on their relative abundance in the metagenomes of investigated samples. "-", not detected.

| Group | MAGs | Comple-teness (%) | Redun-dancy (%) | Genome size (Mb) | GC (%) | Relative abundance (%) | | | | | | | |
|---|---|---|---|---|---|---|---|---|---|---|---|---|---|
| | | | | | | **Larsemann Hills (**Sediment/Soil**)** | | | | | | **South Victoria Land (**soil**)** | |
| | | | | | | **LA1/LS1** | **LA2/LS2** | **LA3/LS3** | **LA4/LS4** | **LA5/LS5** | **LA6/LS6** | **SVL1** | **SVL2** |
| I.1b-AOA | *Nitrososphaera* sp. La5-A1 | 98.06 | 1.94 | 3.563 | 37.9 | -/- | 0.077/- | 0.009/- | 0.013/- | **0.156**/- | -/- | - | - |
| I.1b-AOA | *Nitrosocosmicus* sp. Ls6-A2 | 85.36 | 2.43 | 1.176 | 29.1 | -/- | 0.048/0.008 | 0.012/0.056 | 0.021/- | 0.049/- | -/**0.076** | - | - |
| I.1b-AOA | *Nitrosocosmicus* sp. La5-A2 | 82.52 | 3.4 | 2.473 | 29.6 | -/- | 0.002/- | -/- | -/- | **0.064**/- | -/- | - | - |
| I.1b-AOA | *Nitrosocosmicus* sp. Ls6-A1 | 75.35 | 0.97 | 2.132 | 33.9 | -/- | -/- | -/- | -/- | -/- | -/**0.149** | - | - |
| I.1b-AOA | *Nitrosocosmicus* sp. Ls1-A1 | 52.71 | 5.02 | 1.719 | 29.6 | 0.008/**0.132** | 0.007/- | -/- | -/- | -/- | -/- | - | - |
| Comammox clade B | *Nitrospira* sp. La1-X1 | 96.82 | 2.73 | 4.043 | 56.7 | **0.173**/0.014 | 0.005/- | -/- | -/- | 0.002/- | -/- | - | - |
| Comammox clade B | *Nitrospira* sp. La3-X1 | 91.31 | 4.09 | 3.574 | 56.6 | - | 0.005/- | **0.047**/- | 0.028/0.006 | -/- | 0.027/- | - | - |
| Comammox clade B | *Nitrospira* sp. Ls1-X1 | 84.44 | 1.42 | 3.498 | 55.3 | 0.043/**0.151** | 0.008/- | -/- | -/- | 0.003/- | -/- | - | - |
| Comammox clade B | *Nitrospira* sp. La1-X2 | 77.67 | 7.12 | 5.392 | 55.2 | **0.053**/0.058 | 0.005/- | -/- | -/- | 0.003/- | -/- | - | - |
| NOB lineage IV | *Nitrospira* sp. Ls6-N1 | 76.06 | 5.51 | 2.861 | 50.2 | -/- | -/- | 0.002/- | -/0.011 | -/0.002 | -/**0.108** | - | - |
| NOB lineage IV | *Nitrospira* sp. La5-N1 | 75.72 | 2.49 | 2.703 | 56.0 | -/- | -/- | -/- | -/- | **0.065**/- | -/- | - | - |
| NOB lineage II | *Nitrospira* sp. Ls1-N1 | 73.63 | 3.69 | 2.637 | 56.2 | 0.002/**0.043** | 0.040/- | 0.002/- | -/- | 0.014/- | -/- | - | - |

**Supplementary Table 4.** The gene-based diversity indices for of AOA-, AOB- and comammox *Nitrospira-amoA* and *Nitrospira-nxrB* genes in LA1-LA6 (lake sediments) and LS4 (soil) in Larsemann Hills in coastal East Antarctica. n.a., not available; PD, phylogenetic diversity.

| **Sample** | **AOA/AOB-*amoA*** | | | | | **Comammox *Nitrospira-amoA*** | | | | | ***Nitrospira-nxrB*** | | | | |
|---|---|---|---|---|---|---|---|---|---|---|---|---|---|---|---|
| | Observed OTUs | Shannon | Simpson | Chao | PD | Observed OTUs | Shannon | Simpson | Chao | PD | Observed OTUs | Shannon | Simpson | Chao | PD |
| LA1 | 3/n.a. | 0.1/n.a. | 0.0/n.a. | 3/n.a. | 0.3/n.a. | 36 | 1.6 | 0.5 | 41 | 1.4 | 12 | 1.6 | 0.6 | 12 | 0.5 |
| LA2 | 13/5 | 2.3/2.3 | 0.7/0.8 | 13/15 | 1.2/0.4 | 40 | 2.6 | 0.8 | 40 | 1.6 | 33 | 3.2 | 0.8 | 33 | 1.3 |
| LA3 | 3/5 | 1.0/2.3 | 0.4/0.8 | 3/15 | 0.3/0.5 | 36 | 0.7 | 0.2 | 36 | 1.5 | 24 | 1.3 | 0.4 | 24 | 0.9 |
| LA4 | 6/4 | 1.4/2.0 | 0.6/0.8 | 6/10 | 0.7/0.5 | 41 | 1.5 | 0.4 | 41 | 2.0 | 24 | 1.1 | 0.3 | 26 | 1.0 |
| LA5 | 4/6 | 1.5/2.6 | 0.6/0.8 | 4/21 | 0.4/0.5 | 39 | 2.6 | 0.8 | 47 | 2.0 | 18 | 3.0 | 0.9 | 19 | 0.8 |
| LA6 | n.a./2 | n.a./1.0 | n.a./0.5 | n.a./3 | n.a./0.1 | 35 | 1.0 | 0.2 | 39 | 1.4 | 10 | 1.3 | 0.4 | 10 | 0.5 |
| LS4 | 10/17 | 2.1/4.1 | 0.7/0.9 | 10/153 | 0.9/1.2 | 31 | 1.5 | 0.5 | 32 | 1.3 | 20 | 1.8 | 0.6 | 23 | 0.7 |

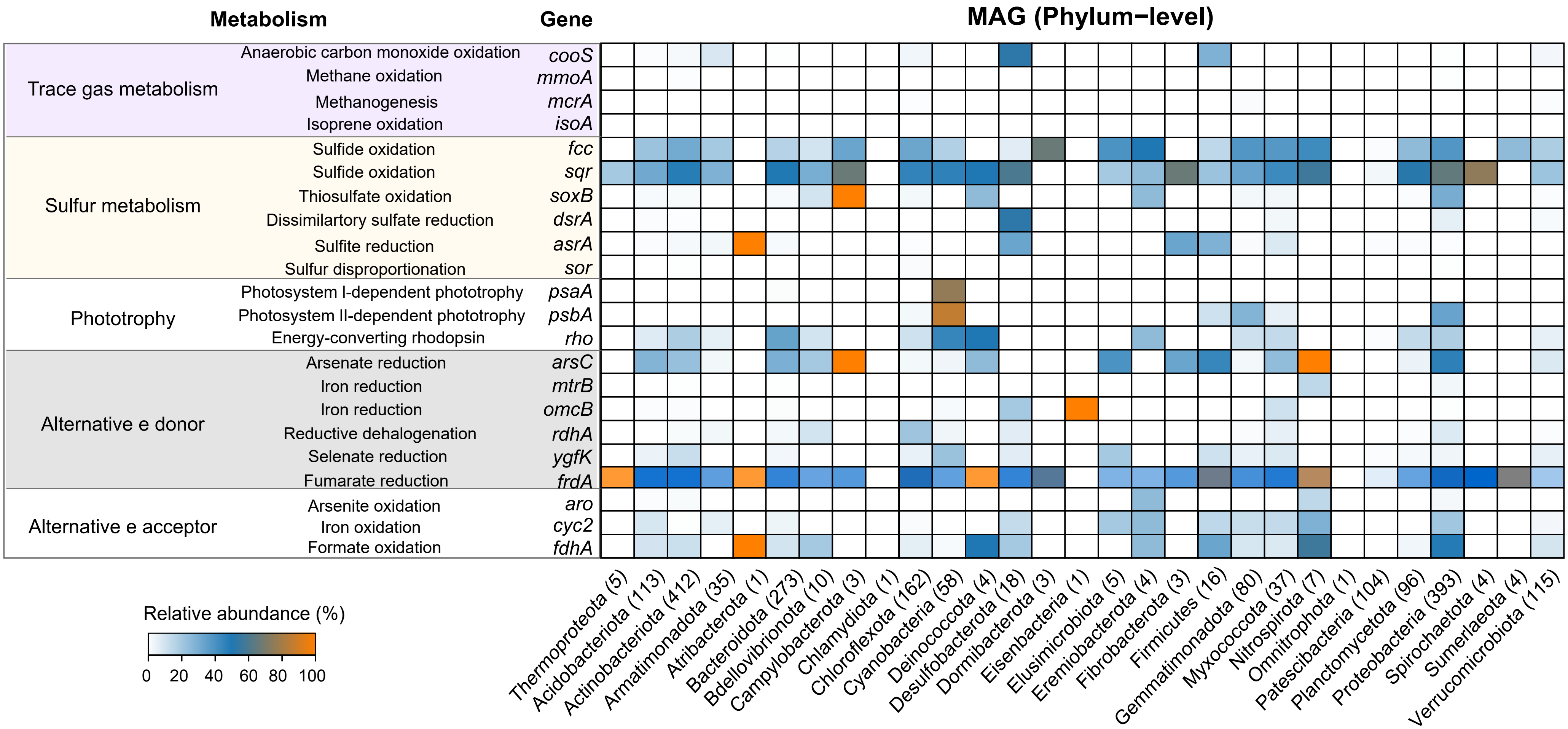


**Supplementary Fig. 1.** Identification and distribution of signature genes encoding enzymes involved in trace gas metabolism, sulfur metabolism, phototrophy, alternative electron donors, and alternative electron acceptors within metagenome-assembled genomes. The heatmap on the right demonstrates the prevalence of these genes across a comprehensive collection of 1968 metagenome-assembled genomes, spanning 29 different phyla.

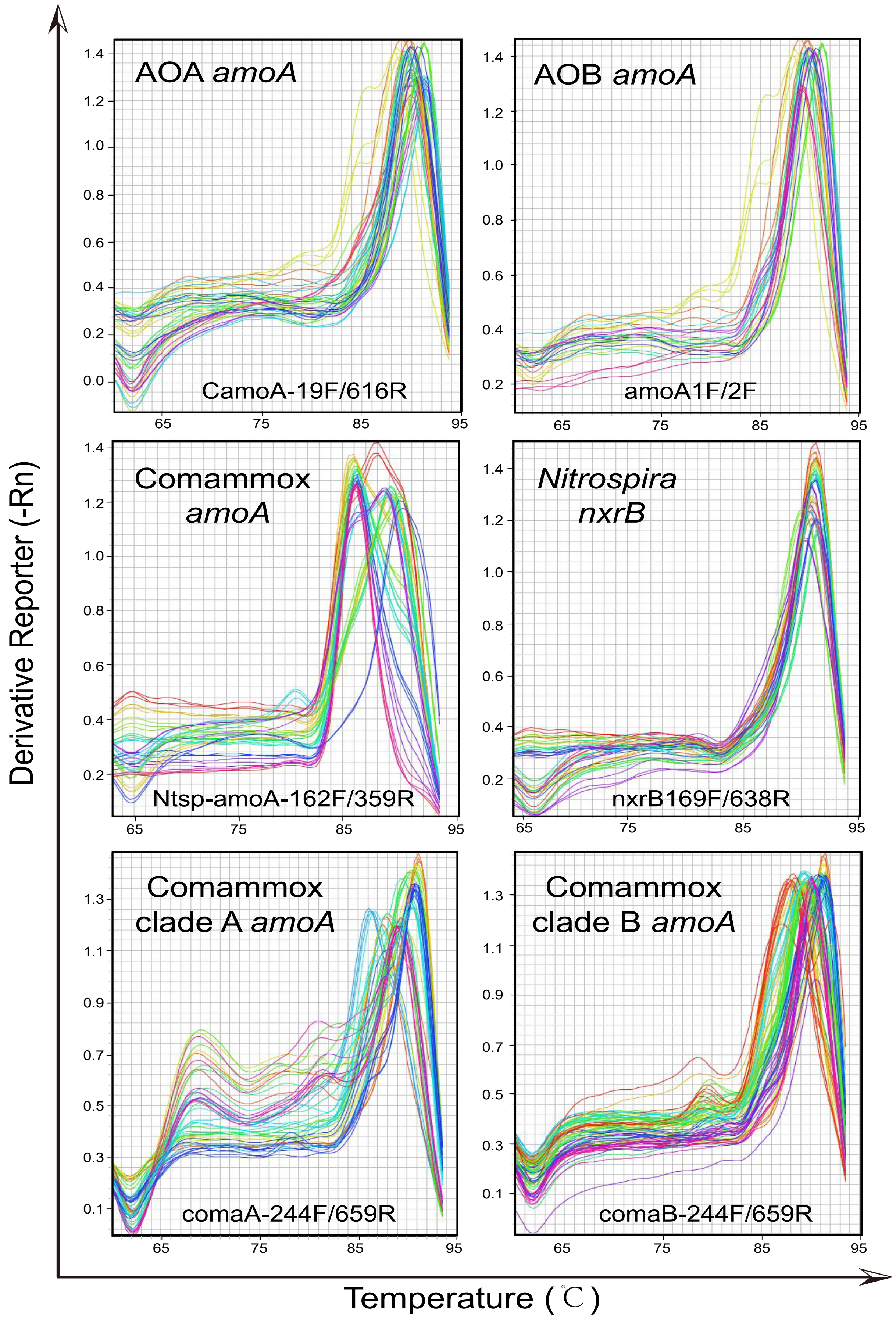


**Supplementary Fig. 2.** Melting curves demonstrating the specificity of various qPCR primers targeting different nitrifying groups including AOA, AOB, comammox and nitrite-oxidizin *Nitrospira*.

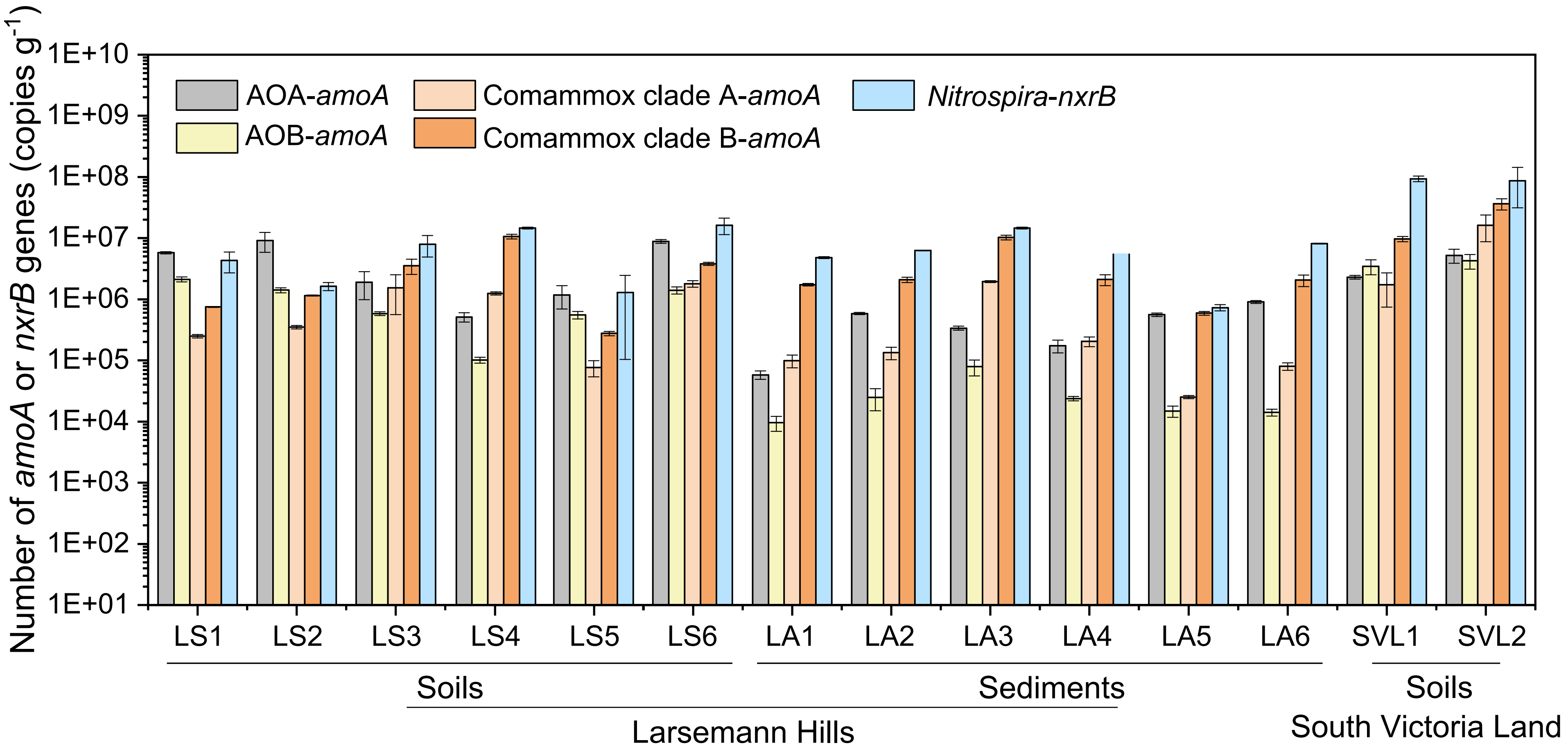


**Supplementary Fig. 3.** Abundance of AOA-, AOB-, comammox-*amoA*, and *Nitrospira*-*nxrB* genes in the samples scrutinized in this study. Comammox clade A and comammox clade B were individually quantified using their respective sets of specific primers listed in Supplementary Table 2.

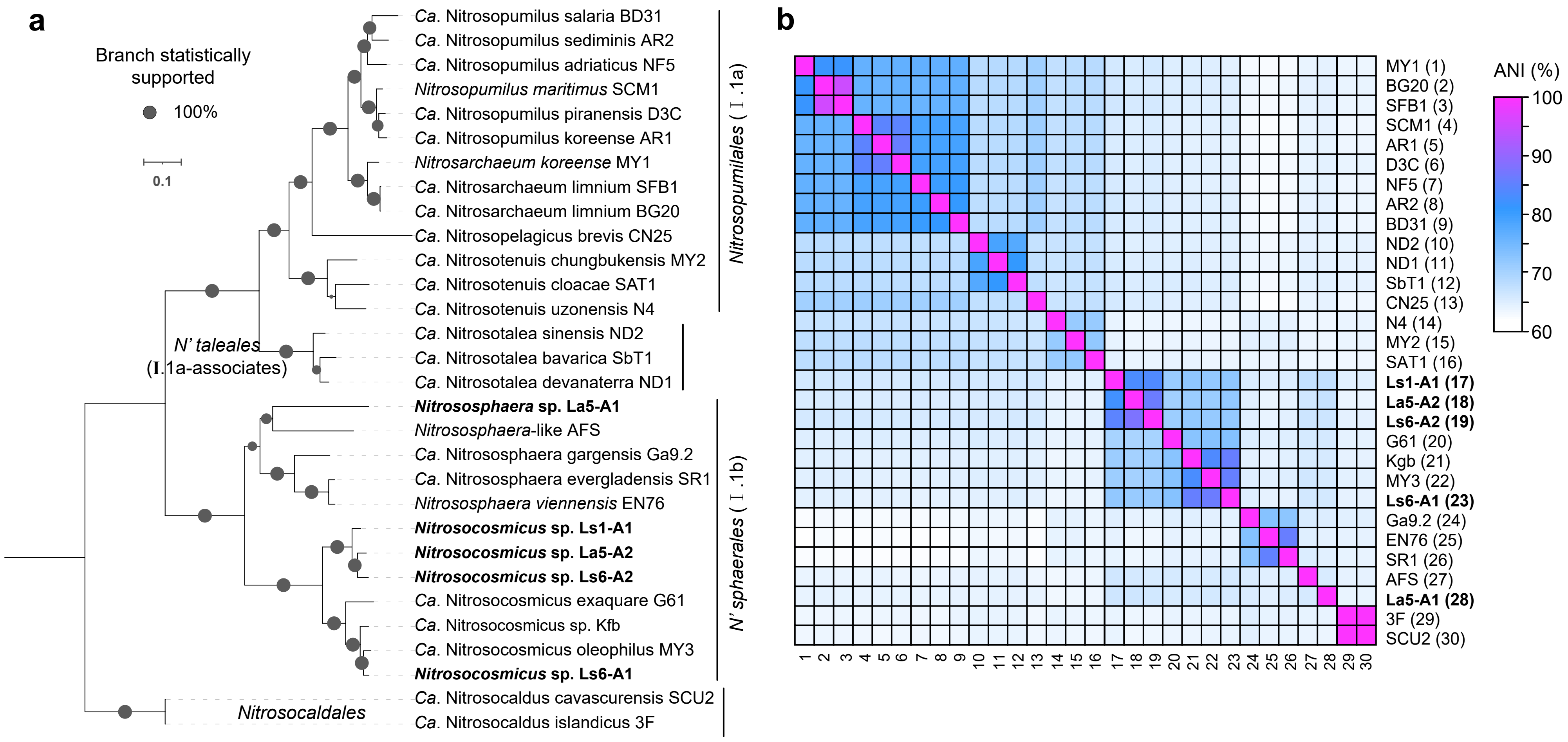


**Supplementary Fig. 4. (a)** Maximum likelihood phylogenomic tree for AOA genomes. **(b)** Average Nucleotide Identity (ANI) analysis on a selected set of AOA genomes from the NCBI database. The AOA metagenome-assembled genomes (MAGs) identified in this study are highlighted in bold.

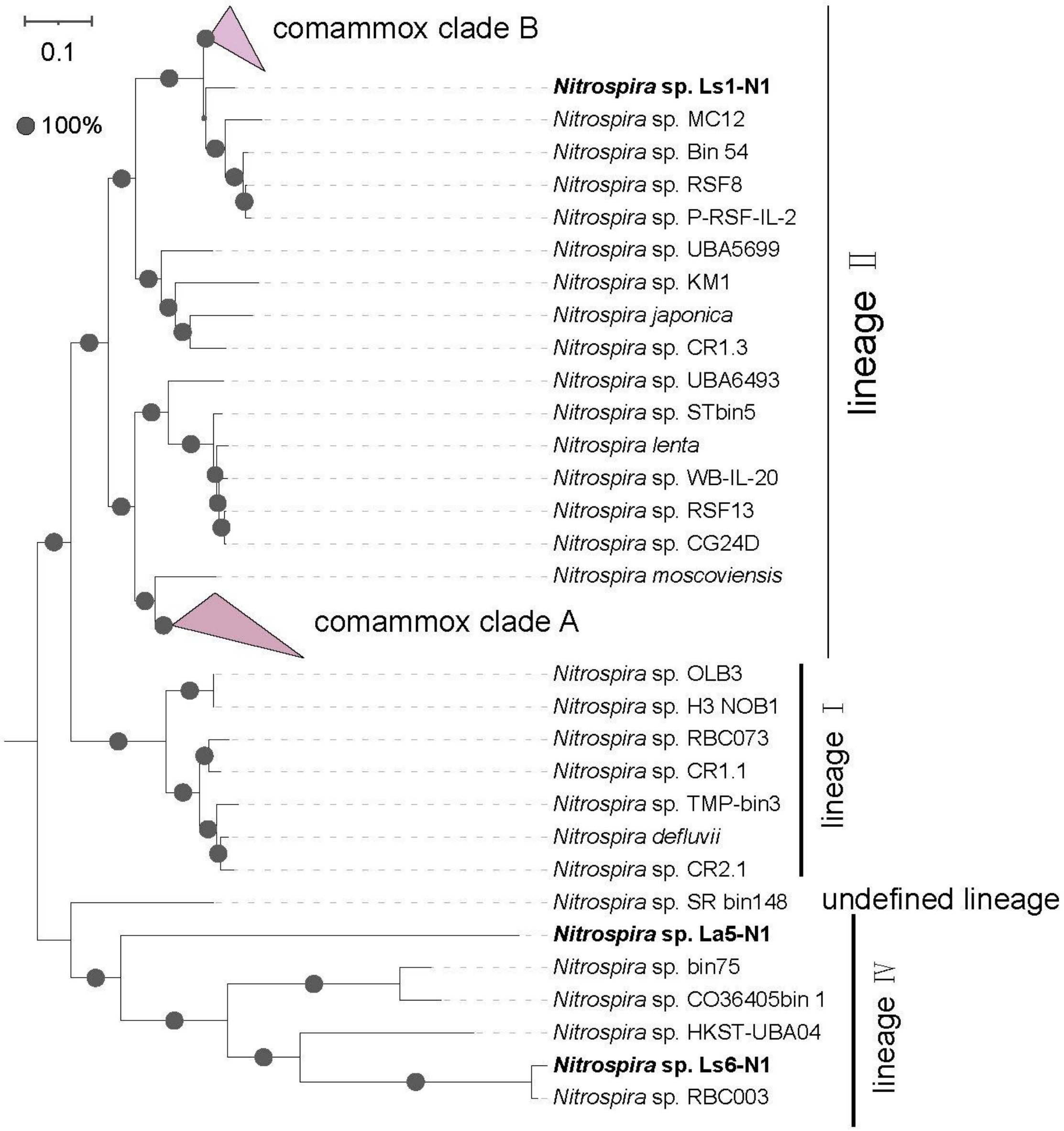


**Supplementary Fig. 5.** Maximum likelihood phylogenomic tree for the genus *Nitrospira*, with a specific focus on the strictly nitrite-oxidizing *Nitrospira*. The *Nitrospira* metagenome-assembled genomes (MAGs) identified in this study are emphasized in bold. The genomes of comammox *Nitrospira* are represented as collapsed branches in the tree.

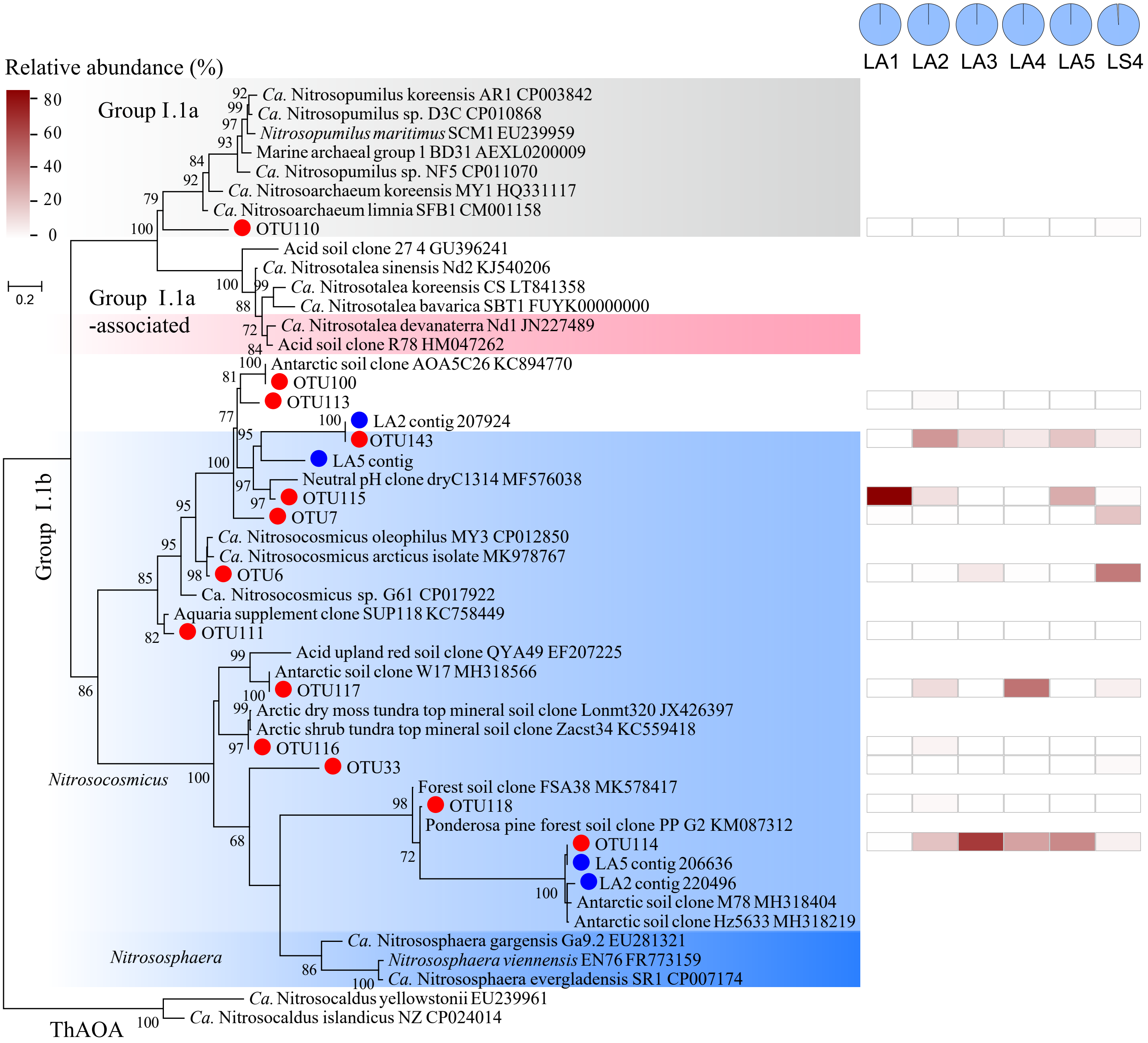


**Supplementary Fig. 6.** Maximum likelihood phylogenetic tree (on the left) and the relative abundance (on the right) of AOA-*amoA* gene sequences in LA1-LA5 (lake sediments) and LS4 (soil) from Larsemann Hills, East Antarctica. The pie charts in the top right corner represent the relative abundance patterns of different AOA groups, marked with the same colors as those in the tree background. The representative OTU sequences obtained in this study are denoted by closed red circles, while sequences from contigs or MAGs are indicated by closed blue circles.

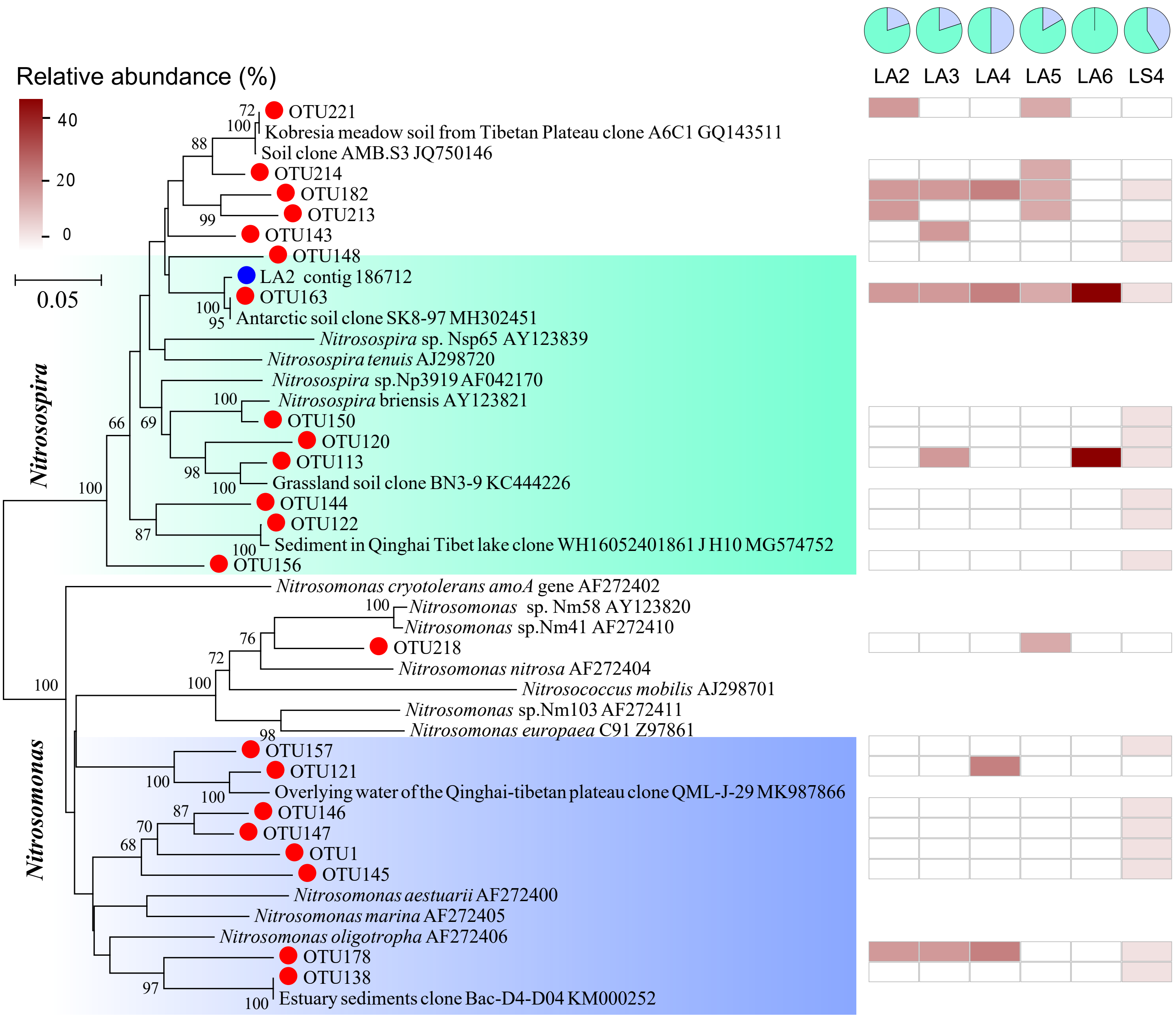


**Supplementary Fig. 7.** Maximum likelihood phylogenetic tree along with the relative abundance of AOB - *amoA* gene sequences in LA2-LA6 (lake sediments) and LS4 (soil) from Larsemann Hills, East Antarctica. The pie charts positioned in the top right corner display the relative abundance patterns of various AOB genera, which are distinguished by the same colors as those in the tree background. The representative OTU sequences obtained in this study are denoted by closed red circles, while sequences from contigs or MAGs are indicated by closed blue circles.

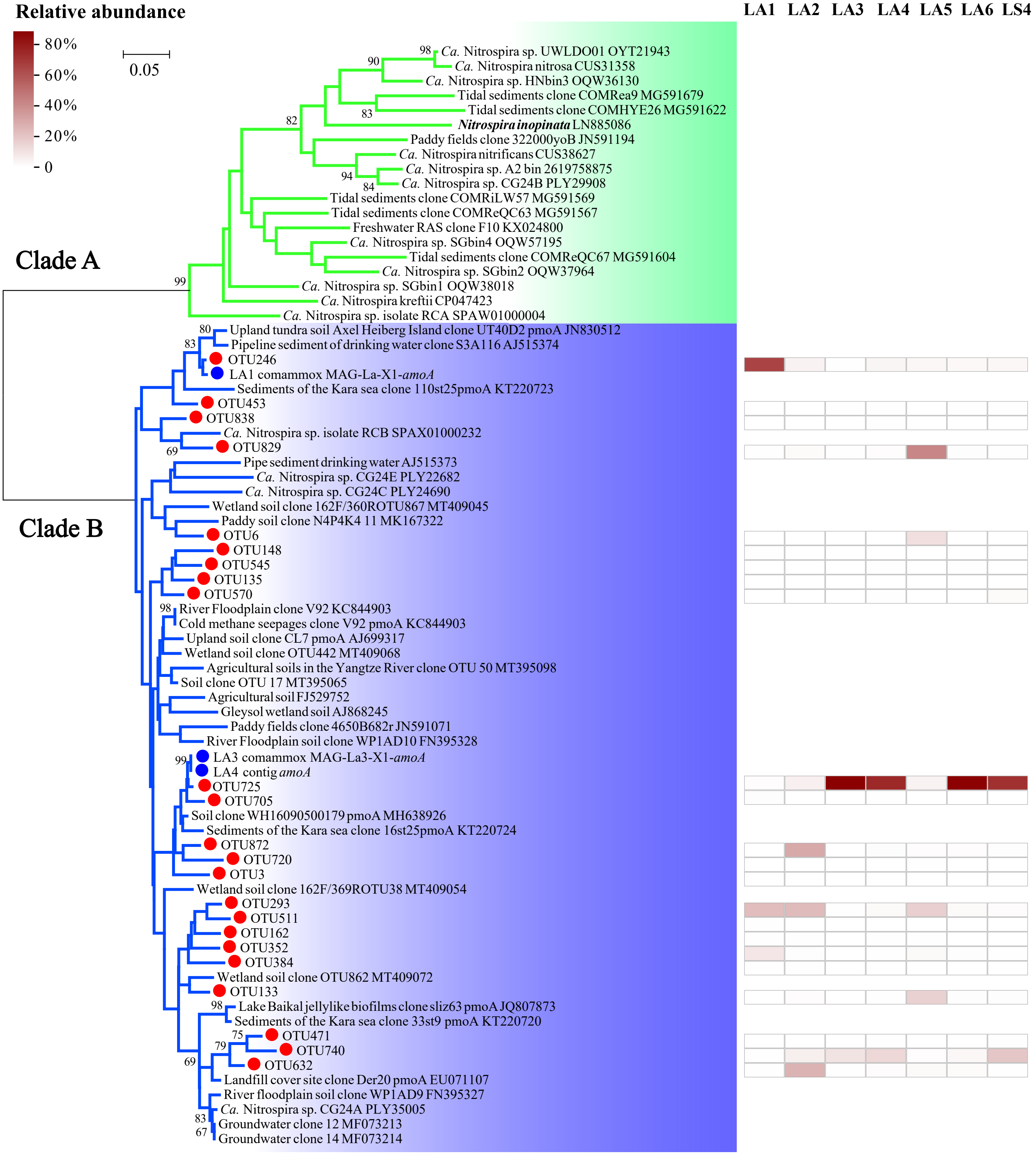


**Supplementary Fig. 8.** Maximum likelihood phylogenetic tree (left) and relative abundance (right) of comammox *Nitrospira-amoA* sequences in LA1-LA6 (lake sediments) and LS4 (soil) from Larsemann Hills, East Antarctica. The representative OTU sequences obtained in this study are denoted by closed red circles, while sequences from contigs or MAGs are indicated by closed blue circles.

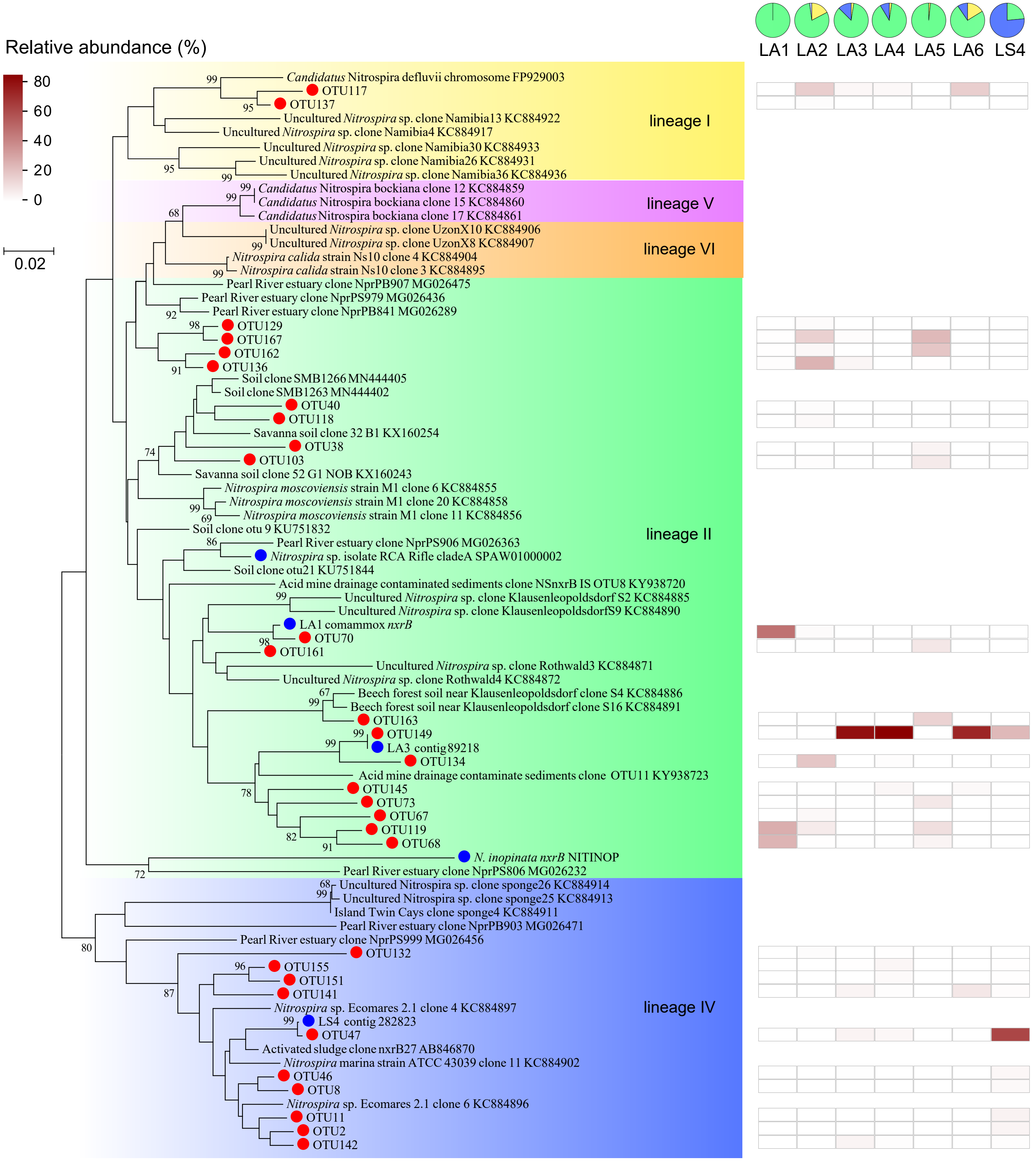


**Supplementary Fig. 9.** Maximum likelihood phylogenetic tree and the relative abundance of *Nitrospira-nxrB* gene sequences in LA1-LA6 (lake sediments) and LS4 (soil) from Larsemann Hills, East Antarctica. The pie charts located in the top right corner demonstrate the relative abundance patterns of different *Nitrospira* lineages, categorized by the same colors as those in the tree background. The representative OTU sequences obtained in this study are denoted by closed red circles, while sequences from contigs or MAGs are indicated by closed blue circles.

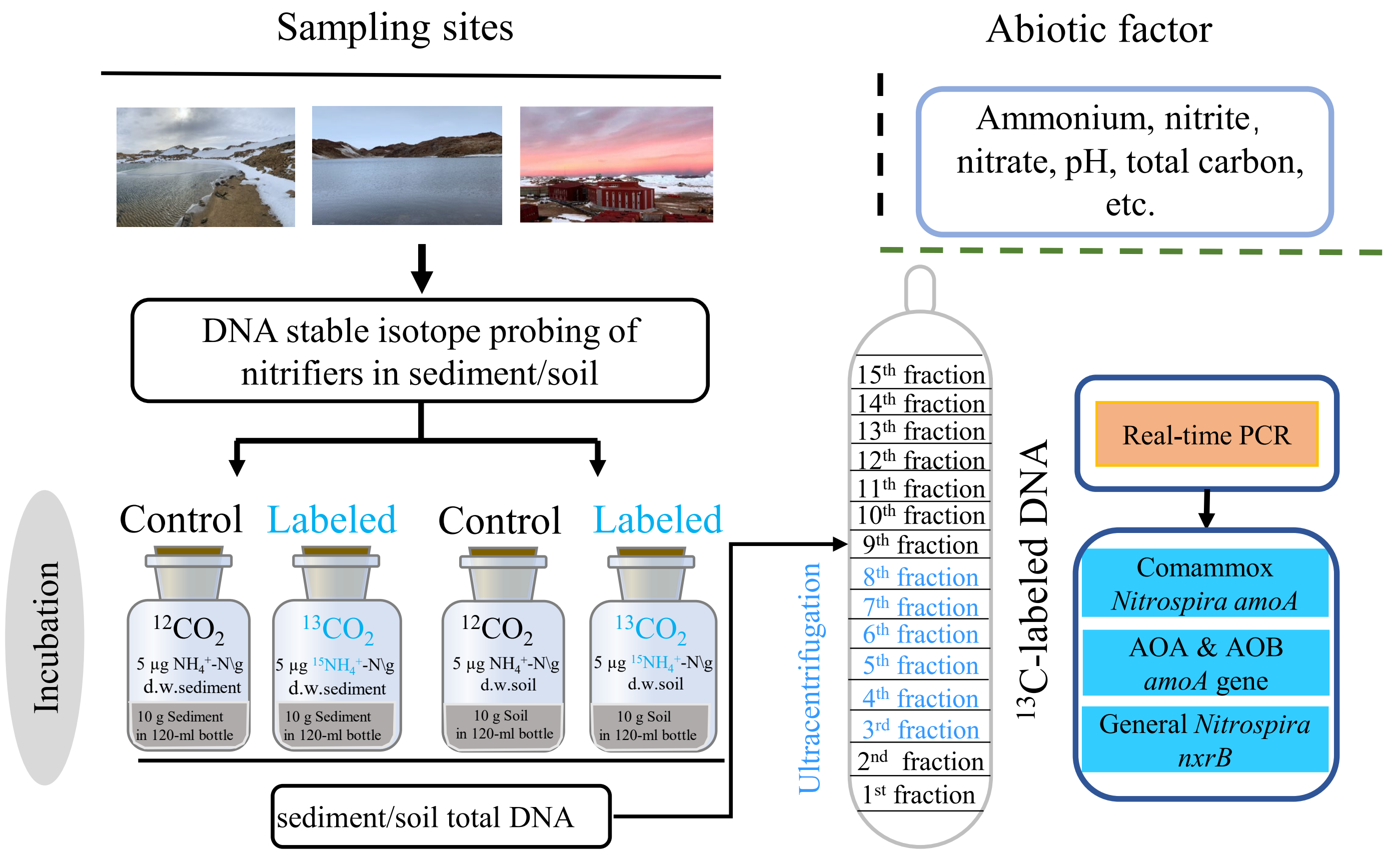


**Supplementary Fig. 10.** Flow chart detailing the DNA-SIP microcosm experiment conducted on two lake sediment samples (LA1 and LA2) and one soil sample (LS4) gathered from Larsemann Hills, East Antarctica.

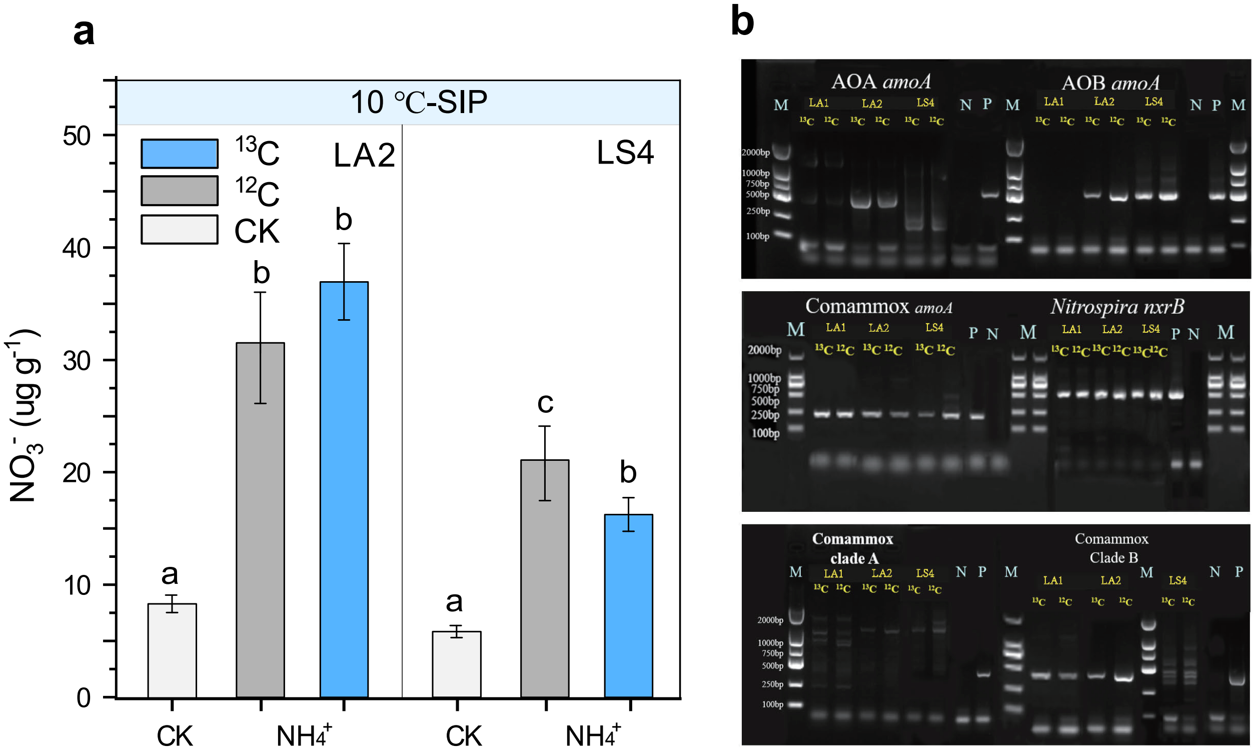


**Supplementary Fig. 11. (a)** Accumulated nitrate concentrations in samples LA2 and LS4 following a 56-day DNA-SIP incubation at a temperature of 10°C, using ammonium as the substrate. The nitrate for LA1 is show in Fig. 4. (b) Agarose gel electrophoresis results of the PCR products for AOA-, AOB-, *comammox-amoA*, and *Nitrospira-nxrB* genes in DNA extracted from the LA1 and LA2 sediment and LS4 soil after DNA-SIP microcosm incubations at 10 °C.

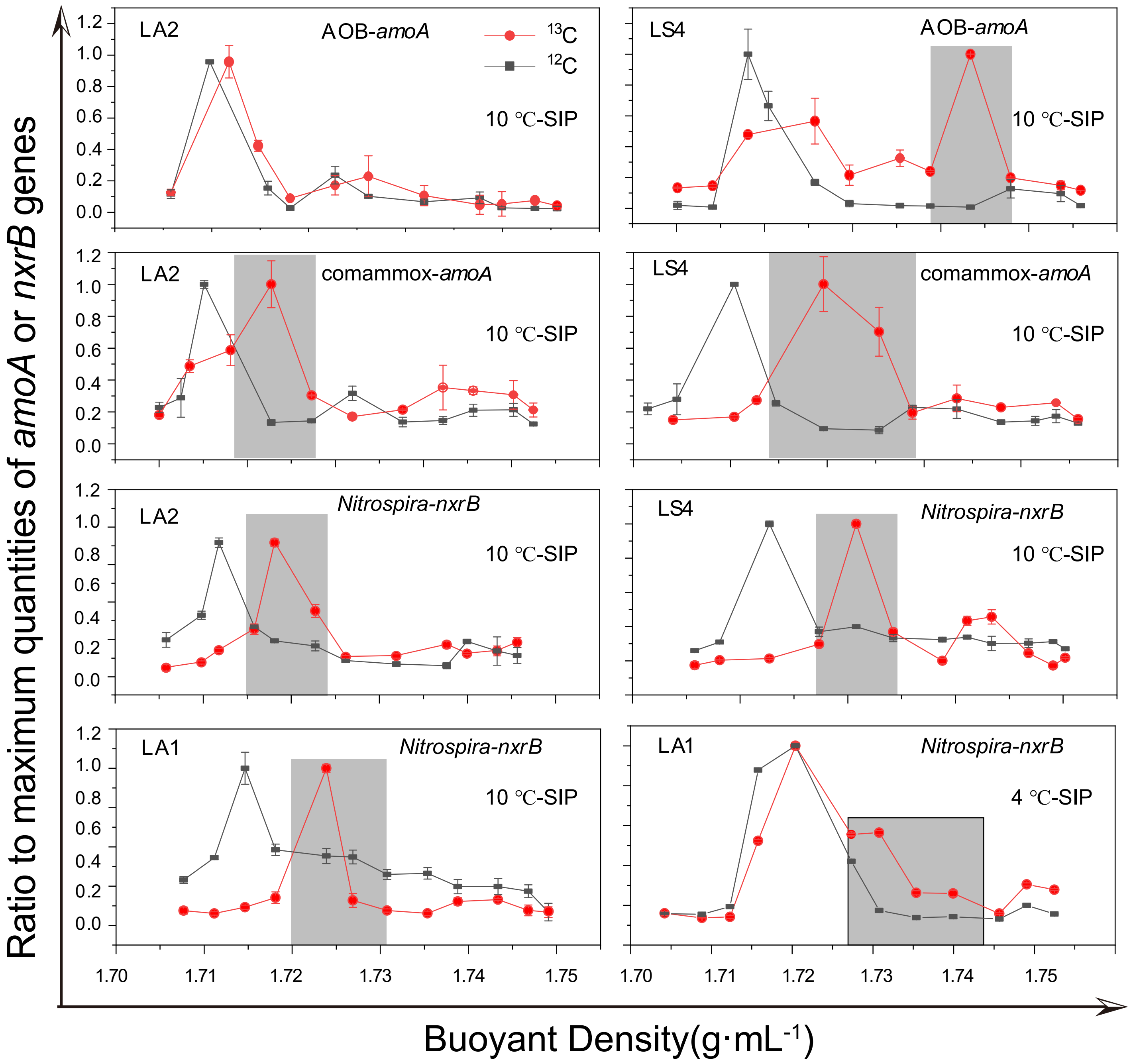


**Supplementary Fig. 12**. Evidence of nitrification activity of AOB, comammox *Nitrospira* and total *Nitrospira.* DNA-SIP-derived quantitative distribution and relative abundance of AOB- and comammox *Nitrospira -amoA* and *Nitrospira-nxrB* genes in $^{13}CO_2$ and $^{12}CO_2$ treated microcosms containing sediment from LA2 (lake sediments), and LS4 (soil) after 56-day incubation at 10 °C or 4 °C, respectively. The outcome for *Nitrospira-nxrB* in LA1 is also included (see bottom), while the labeling for its comammox *Nitrospira-amoA* is depicted in Fig. 4. Error bars indicate standard deviations (n = 3).

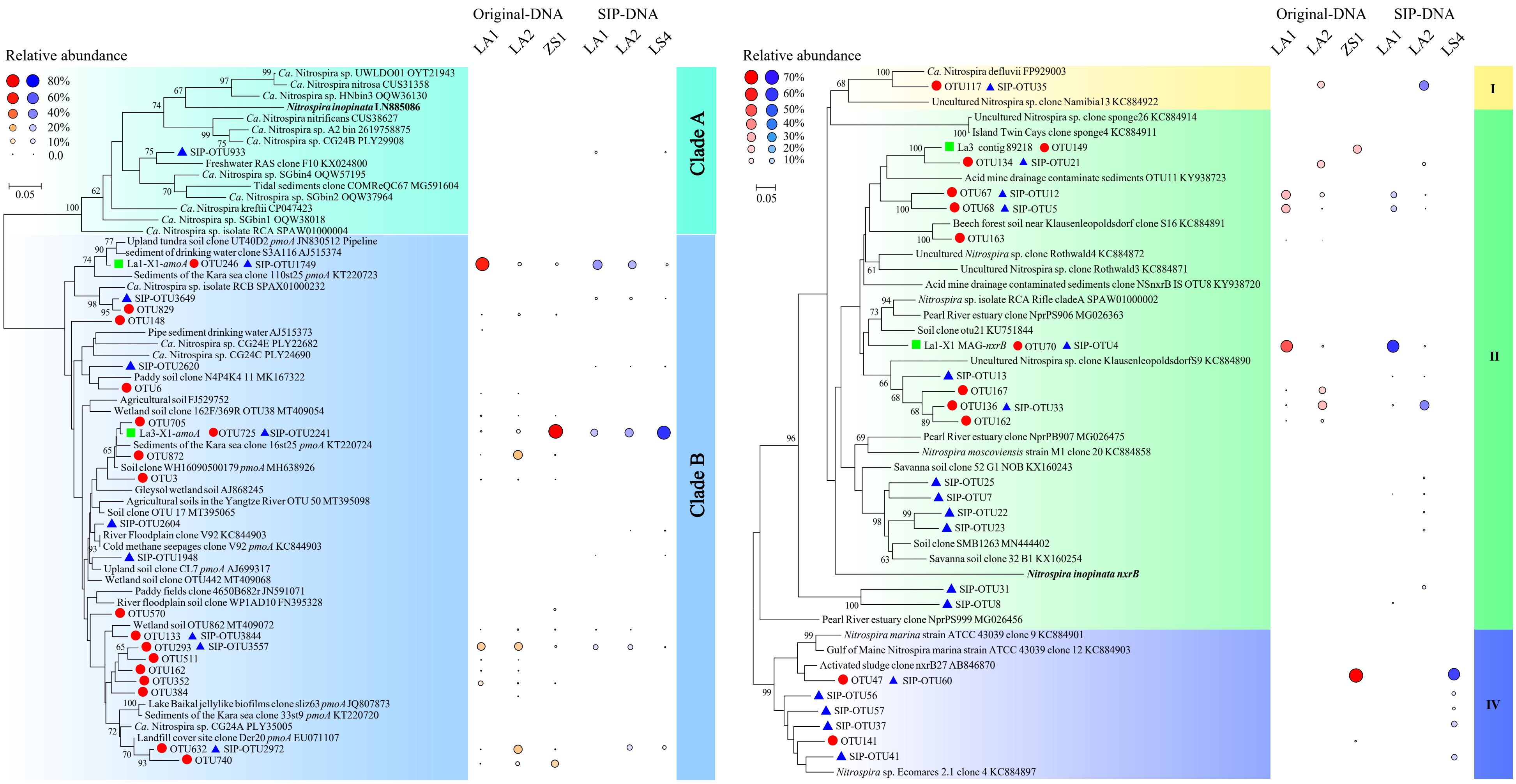


**Supplementary Fig. 13. Comparison of the diversity and relative abundance of *Nitrospira* from original- and 10 °C incubation SIP-DNA.** Maximum-likelihood phylogenetic trees, relative abundances of OTUs in original sample DNA (closed red circle) and heavy-fraction DNA (closed blue triangle) from $^{13}CO_2$-SIP incubations of **(a)** comammox *Nitrospira-amoA* gene sequences and **(b)** *Nitrospira*-affiliated *nxrB* gene sequences. *AmoA* and *nxrB* genes from the retrieved contigs or MAGs in this study are indicated in green square.
